# Tetraose steroidal glycoalkaloids from potato provide resistance against *Alternaria solani* and Colorado potato beetle

**DOI:** 10.1101/2022.09.28.509958

**Authors:** Pieter J. Wolters, Doret Wouters, Yury M. Tikunov, Shimlal Ayilalath, Linda P. Kodde, Miriam Strijker, Lotte Caarls, Richard G. F. Visser, Vivianne G. A. A. Vleeshouwers

**Affiliations:** Plant Breeding, Wageningen University and Research, P.O. Box 386, 6700, AJ, Wageningen, The Netherlands

## Abstract

Plants with innate disease and pest resistance can contribute to more sustainable agriculture. Natural defence compounds produced by plants have the potential to provide a general protective effect against pathogens and pests, but they are not a primary target in resistance breeding. Here, we identified a wild relative of potato, *Solanum commersonii*, that provides us with unique insight in the role of glycoalkaloids in plant immunity. We cloned two atypical resistance genes that provide resistance to *Alternaria solani* and Colorado potato beetle through the production of tetraose steroidal glycoalkaloids. Moreover, we provide *in vitro* evidence to show that these compounds have potential against a range of different (potato pathogenic) fungi. This research links structural variation in steroidal glycoalkaloids to resistance against potato diseases and pests. Further research on the biosynthesis of plant defence compounds in different tissues, their toxicity, and the mechanisms for detoxification, can aid the effective use of such compounds to improve sustainability of our food production.

## Introduction

Worldwide, up to 20-40% of agricultural crop production is lost due to plant diseases and pests (1). Many crops have become heavily dependent on the use of pesticides, but this is unsustainable as these can negatively affect the environment and their use can lead to development of pesticide resistance (2–7). The European Union’s ‘Farm to Fork Strategy’ aims to half pesticide use and risk by 2030 (8), a massive challenge that illustrates the urgent need for alternative disease control methods.

Wild relatives of crop species are promising sources of natural disease resistance (9–12). Monogenic resistance caused by dominant resistance (*R*) genes, typically caused by immune receptors that belong to the nucleotide-binding leucine-rich repeat (NLR) class, is successfully employed by plant breeders to develop varieties with strong qualitative disease resistance. However, this type of resistance is usually restricted to a limited range of pathogens (13, 14) and it is often not very durable.

More robust resistance can be obtained by combining NLRs with different recognition specificities (15–18), or by including pattern recognition receptors (PRRs), which recognize conserved (microbe-or pathogen-derived) molecular patterns. Recent reports show that PRRs and NLRs cooperate to provide disease resistance (19–21). Alternatively, susceptibility (*S*) genes provide recessive resistance that can be both broad-spectrum and durable (22–24). Unfortunately, their recessive nature complicates the use of *S* genes in conventional breeding of autopolyploids and many mutated *S* genes come with pleiotropic effects.

Besides the defences mentioned above, most plants produce specialised metabolites with antimicrobial or anti-insect activity, either constitutively (phytoanticipins) or in response to pathogen attack or herbivory (phytoalexins) (25). These natural defence compounds are derived from a wide range of building blocks, leading to a large structural diversity throughout the plant kingdom (26, 27). Specific classes of compounds can be found in different plant families, e.g. glucosinolates are typically found in Brassicaceae and benzoxyzinoids are widely distributed among Poaceae, with further chemical diversification within plant families (28, 29). Examples in various pathosystems show that the capacity for detoxification of plant defence compounds is important for pathogenicity, especially for necrotrophic pathogens which encounter toxic plant metabolites as a consequence of their lifestyle (30, 31). Similarly, plant defence compounds play an important role against herbivorous insects, which have evolved various mechanisms to deal with toxic plant compounds (32–34).

While plant immune receptors can offer resistance against a restricted range of pathogens, plant defence compound have the potential to provide a more general protection, depending on their mode of action. Plants from the nightshade family (Solanaceae) produce saponins that are characterised by a steroidal alkaloid aglycone, linked to a variable oligosaccharide chain (steroidal glycoalkaloids – SGAs) (35, 36). The protective effect of SGAs stems from their ability to interact with membrane sterols, disrupting the cell integrity from target organisms (37–40). In addition, they can act on the nervous system of pests and herbivores through their inhibitory effect on cholinesterase enzymes (41–43). As a consequence, SGAs can have both antimicrobial and anti-insect activity (44–54).

Early blight is an important disease of tomato and potato that is caused by the necrotrophic fungal pathogen *Alternaria solani* (55–57). In a previous study, we found a wild potato species, *Solanum commersonii*, with strong resistance against potato pathogenic *Alternaria* isolates and species from a number of different locations (58). We showed that resistance is likely caused by a single dominant locus and that it can be introgressed in cultivated potato (58). Resistance to necrotrophs is usually considered to be a complex, polygenic trait, or recessively inherited according to the *inverse gene-for-gene* model (59–64). It therefore surprised us to find a qualitative dominant resistance against early blight in *S. commersonii* (58).

In this study, we explored different accessions of *S. commersonii* and *S. malmeanum* (previously also referred to as *S. commersonii* subsp. *malmeanum* (65)) and developed a population that segregates for resistance to early blight. Using a Bulked Segregant RNA-Seq (BSR-Seq) approach (66), we mapped the resistance locus to the top of chromosome 12 of potato. We sequenced the genome of the resistant parent of the population and identified two glycosyltransferases that can provide resistance to susceptible *S. commersonii*. We show that the resistance is based on the production of tetraose SGAs and provide *in vitro* evidence to show that they can be effective against other fungi besides *A. solani*. As SGAs can be involved in resistance against insects, we also tested if they can protect against Colorado potato beetle (CPB). Combined, our results show that the tetraose SGAs from *S. commersonii* have potential to provide resistance against a range of potato pathogens and pests.

## Results

### Early blight resistance maps to chromosome 12 of potato

To find suitable parents for a mapping study targeting early blight resistance, we performed a disease screen with *A. solani* isolate altNL03003 (67) on 13 different accessions encompassing 37 genotypes of *S. commersonii* and *S. malmeanum* (S1 Table). The screen showed clear differences in resistance phenotypes between and within accessions (Fig 1A). Roughly half of the genotypes were highly resistant (lesion diameters < 3 mm indicate that the lesions are not expanding beyond the size of the inoculation droplet) and the other half was susceptible (displaying expanding lesions), with only a few intermediate genotypes. CGN18024 is an example of an accession that segregates for resistance, with CGN18024_1 showing strong resistance and CGN18024_3 showing clear susceptibility (Fig 1B). The fact that individual accessions can display such clear segregation for resistance suggests that resistance is caused by a single gene or locus. Because of its clear segregation, *S. commersonii* accession CGN18024 was selected for further studies.

**Fig 1.**
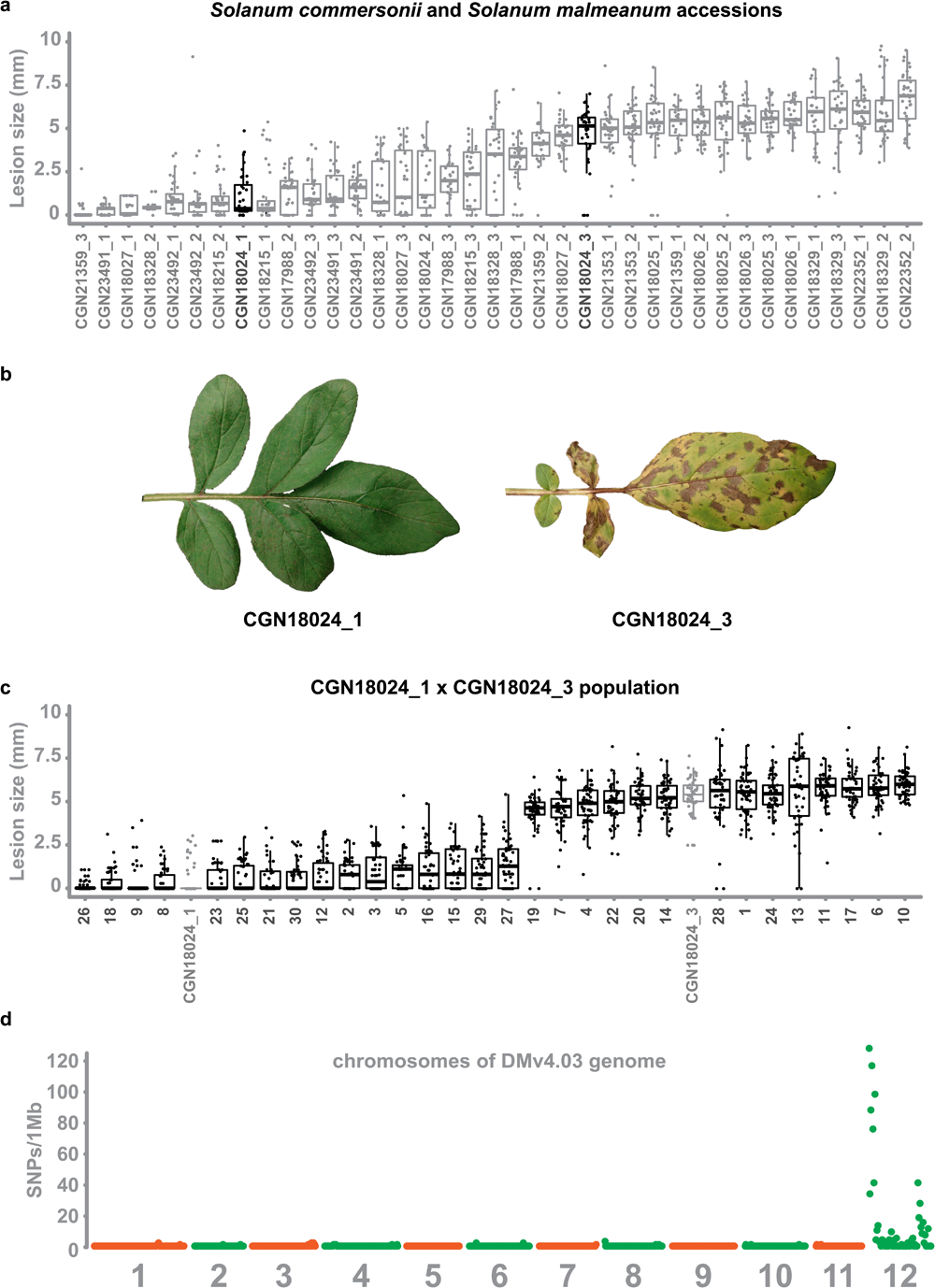
Early blight resistance maps to chromosome 12 of potato. **A.** 2-3 genotypes of 13 different accessions of *S. commersonii* and *S. malmeanum* were inoculated with *A. solani* altNL03003. 3 plants of each genotype were tested and 3 leaves per plants were inoculated with 6 10 µl droplets with spore suspension. Lesion diameters were measured 5 days post inoculation and visualised using boxplots, with horizonal lines indicating median values and individual measurements plotted on top. Non-expanding lesions (<3 mm) indicate resistance and expanding lesions indicate susceptibility. Some accessions segregate for resistance. **B.** Accession CGN18024 is an example of an accession that segregates for resistance to *A. solani*, with CGN18024_1 displaying resistance and CGN18024_3 displaying susceptibility at 5 days after spray-inoculation. **C.** Progeny from CGN18024_1 x CGN18024_3 was inoculated with *A. solani*. 3 plants of each genotype were tested and 3 leaves per plants were inoculated with 6 10 µl droplets with spore suspension each. Lesion diameters were measured 5 days post inoculation. 16 progeny genotypes are resistant (with lesion diameters < 2-3 mm) and 14 are susceptible (with expanding lesions). This corresponds to a 1:1 segregation ratio (Χ^2^ (1, N = 30) = 0.133, ρ= 0.72). **D.** SNPs derived from a BSRseq analysis using bulks of susceptible and resistant progeny were plotted in 1 Mb windows over the 12 chromosomes of the potato DMv4.03 genome (68) They are almost exclusively located on chromosome 12.

Disease tests with an *A. solani* isolate from the US (ConR1H) and a recent Dutch isolate from the Netherlands (altNL21001) confirm that the resistance of CGN18024_1 is effective against additional *A. solani* isolates (S1 Fig). To further study the genetics underlying resistance to early blight, we crossed resistant CGN18024_1 with susceptible CGN18024_3. Thirty progeny genotypes were sown out and tested with *A. solani* isolate altNL03003. We identified 14 susceptible genotypes and 16 resistant genotypes, with no intermediate phenotypes in the population (Fig 1C). This segregation supports a 1:1 ratio (Χ (1, N = 30) = 0.133, p=.72), which confirms that resistance to early blight is likely caused by a single dominant locus in *S. commersonii*.

To genetically localize the resistance, we isolated RNA from each progeny genotype and the parents of the population and proceeded with a BSR-Seq analysis (66). RNA from resistant and susceptible progeny genotypes were pooled in separate bulks and sequenced next to RNA from the parents on the Illumina sequencing platform (PE150). Reads were mapped to the DMv4.03 (68) and Solyntus potato genomes (69). To find putative SNPs linked to resistance, we filtered for SNPs that follow the same segregation as resistance (heterozygous in resistant parent CGN18024_1 and the resistant bulk, but absent or homozygous in susceptible parent and susceptible bulk). The resulting SNPs localize almost exclusively on chromosome 12 of the DM and Solyntus genomes, with most of them located at the top of the chromosome (Fig 1D). We used a selection of SNPs distributed over chromosome 12 as high-resolution melt (HRM) markers to genotype the BSR-Seq population. This rough mapping proves that the locus for early blight resistance resides in a region of 3 Mb at the top of chromosome 12 (S2 Fig).

### Improved genome assembly ***of S. commersonii***

A genome sequence of *S. commersonii* is already available (70), but we do not know if the sequenced genotype is resistant to *A. solani*. To help the identification of additional markers and to explore the resistance locus of a genotype with confirmed resistance, we sequenced the genome of resistant parent CGN18024_1. High-molecular-weight (HMW) genomic DNA (gDNA) from CGN18024_1 was used for sequencing using Oxford Nanopore Technology (ONT) on a GridION X5 platform and for sequencing using DNA Nanoball (DNB) technology at the Beijing Genomics Institute (BGI) to a depth of approximately 30X. We used the ONT reads for the initial assembly and the shorter, more accurate, DNBseq reads to polish the final sequence. The resulting assembly has a size of 737 Mb, which is close to the size of the previously published genome of *S. commersonii* (730 Mb) (70). N50 scores and Benchmarking Universal Single-Copy Orthologs (BUSCO) score indicate a highly complete and contiguous genome assembly of *S. commersonii* (Table 1).

**Table 1.** Genome assembly metrics of *S. commersonii* cmm1t (70) and CGN18024_1.

| Genome | CMM1t<sup>a)</sup> | CGN18024_1 |
| --- | --- | --- |
| Total size (Mb) | 730 | 737 |
| Contig number | 278460 | 637 |
| Largest contig (Mb) | 0,17 | 21,2 |
| N50 (Mb) | 0,007 | 4,02 |
| Complete BUSCO (%) | 81,9 | 95,7 |

<sup>a)</sup> Aversano et al (2015)

### Identification of two glycosyltransferase resistance genes

To identify candidate genes that can explain the resistance of *S. commersonii*, it was necessary to further reduce the mapping interval. By aligning the ONT reads to the CGN18024_1 genome assembly, we could identify new polymorphisms that we converted to additional PCR markers (S3-6 Figs). We performed a recombinant screen of approximately 3000 genotypes from the population to fine-map the resistance region to a window of 20 kb (S7 and S8 Figs).

We inferred that the resistance locus is heterozygous in CGN18024_1 from the segregation in the mapping population. We used polymorphisms in the resistance region to separate and compare the ONT sequencing reads from the resistant and susceptible haplotype. This comparison showed a major difference between the two haplotypes. The susceptible haplotype contains a small insertion of 3.7 kb inside a larger region of 7.3 kb. The larger region is duplicated in the resistant haplotype (Fig 2A). As a result, the resistance region of the resistant haplotype is 27 kb, 7 kb larger than the corresponding region of the susceptible haplotype (20 kb).

**Fig 2.**
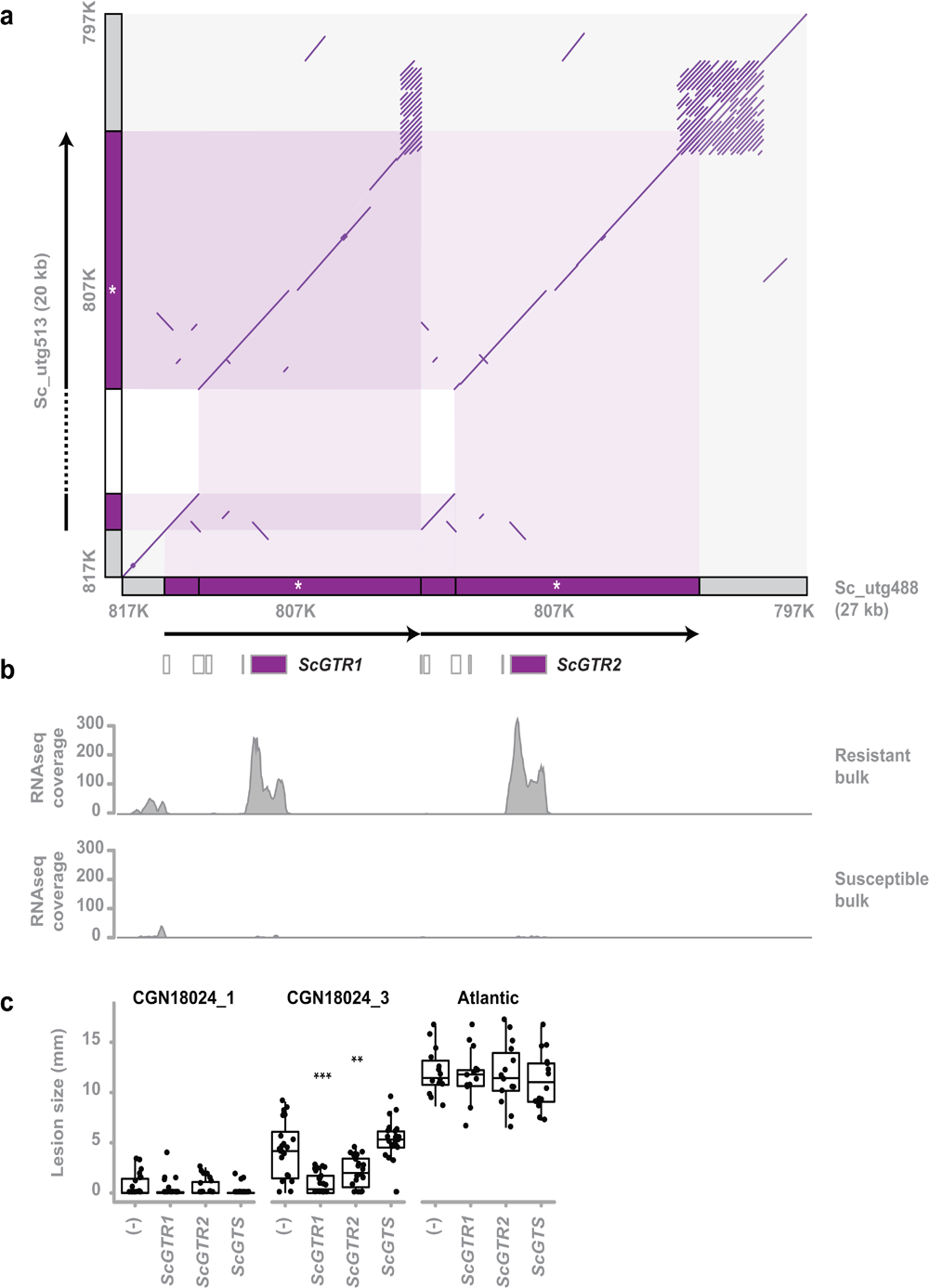
Identification of two glycosyltransferase resistance genes. **A.** Comparison of the susceptible and resistant haplotype of the *Solanum commersonii* CGN18024_1 resistance region (delimited by markers 817K and 797K) in a comparative dot plot shows a rearrangement. Locations of markers used to map the resistance region are indicated in grey along the x-and y-axis. The duplicated region of the resistant haplotype contains marker 807K (white asterisk) and two predicted glycosyltransferase genes (*ScGTR1* and *ScGTR2*). Several short ORFs with homology to glycosyltransferase genes that were predicted in the resistance region are indicated by white boxes, but *ScGTR1* and *ScGTR2* are the only full-length genes. As a result of the rearrangement, the resistance region of the resistant haplotype (27 kb) is 7 kb larger than the corresponding region of the susceptible haplotype (20 kb). **B.** Alignment of RNAseq reads from the BSRSeq analysis shows that *ScGTR1* and *ScGTR2* are expressed in bulks of resistant progeny, but not in bulks of susceptible progeny. **C.** *S. tuberosum*cv. ‘Atlantic’, *S. commersonii* CGN18024_1 and CGN18024_3 were agroinfiltrated with expression constructs for *ScGTR1* and *ScGTR2*, *ScGTS* and empty vector (-). *A. solani* is inoculated 2 days after agroinfiltration and lesion diameters are measured 5 days after inoculation. Lesion sizes were visualised with boxplots, with horizonal lines indicating median values and individual measurements plotted on top. Agroinfiltration with expression constructs for *ScGTR1* and *ScGTR2* results in a significant (Welch’s Two Sample t-test, \*\**P* < 0.01, \*\*\**P* < 0.001) reduction of lesion sizes produced by *Alternaria solani* altNL03003 in *S. commersoni i*CGN18024_3, but not in *S. tuberosum* cv. ‘Atlantic’.

Two genes coding for putative glycosyltransferases (GTs) are located within the rearrangement of the resistant haplotype. The corresponding allele from the susceptible haplotype contains a frameshift mutation, leading to a truncated protein (S9 Fig). Several other short ORFs with homology to glycosyltransferases were predicted in the resistant haplotype, but *ScGTR1* (*S. commersonii* glycosyltransferase linked to resistance 1) and *ScGTR2* are the only full-length genes in the region.

Reads from the BSR-Seq experiment show that both genes are expressed in bulks of resistant progeny and not in susceptible progeny (Fig 2B), suggesting a putative role for these genes in causing resistance. *ScGTR1* and *ScGTR2* are homologous genes with a high similarity (the predicted proteins that they encode share 97% amino acid identity). We compared the predicted amino acid sequences with previously characterised GTs (71–78) and found that they share some similarity with GTs with a role in zeatin biosynthesis (79–81) and with GAME17, an enzyme involved in biosynthesis of the SGA α-tomatine typically found in tomato (76) (S10 Fig, S2 Table).

To test whether the identified candidate genes are indeed involved in resistance, we transiently expressed both alleles of the resistant haplotype (*ScGTR1* and *ScGTR2*) as well as the corresponding allele from the susceptible haplotype (*ScGTS*), in leaves of resistant CGN18024_1 and susceptible CGN18024_3 and *S. tuberosum* cultivar Atlantic, using agroinfiltration (82). Following agroinfiltration, the infiltrated areas were drop-inoculated with a spore suspension of *A. solani* altNL03003. Transient expression of *ScGTR1* as well as *ScGTR2* significantly reduced the size of the *A. solani* lesions in susceptible CGN18024_3, compared to *ScGTS* and the empty vector control. Resistant CGN18024_1 remained resistant, whereas susceptible Atlantic remained susceptible regardless of the treatment (Fig 2C). We conclude that both *ScGTR1* and *ScGTR2* can affect resistance in susceptible *S. commersonii* CGN18024_3, but not in *S. tuberosum* cv. Atlantic.

### Leaf compounds from resistant ***S. commersonii*** inhibit growth of diverse fungi, including pathogens of potato

Glycosyltransferases are ubiquitous enzymes that catalyse the transfer of saccharides to a range of different substrates. To test if resistance of *S. commersonii* to *A. solani* can be explained by a host-specific defence compound, we performed a growth inhibition assay using crude leaf extract from resistant and susceptible *S. commersonii*. Leaf material was added to PDA plates to equal 5% w/v and autoclaved (at 121 °C) or semi-sterilised at 60 °C. Interestingly, leaf material from resistant CGN18024_1 strongly inhibited growth of *A. solani* isolate altNL03003, while we did not see any growth inhibition on plates containing leaves from susceptible CGN18024_3 (Fig 3A). Remarkably, ample contamination with diverse fungi appeared after a few days on the plates containing semi-sterilised leaves from susceptible *S. commersonii* but not on plates with leaves from CGN18024_1 (Fig 3A). Thus, leaves from CGN18024_1 contain compounds that can inhibit growth of a variety of fungi besides *A. solani*.

**Fig 3.**
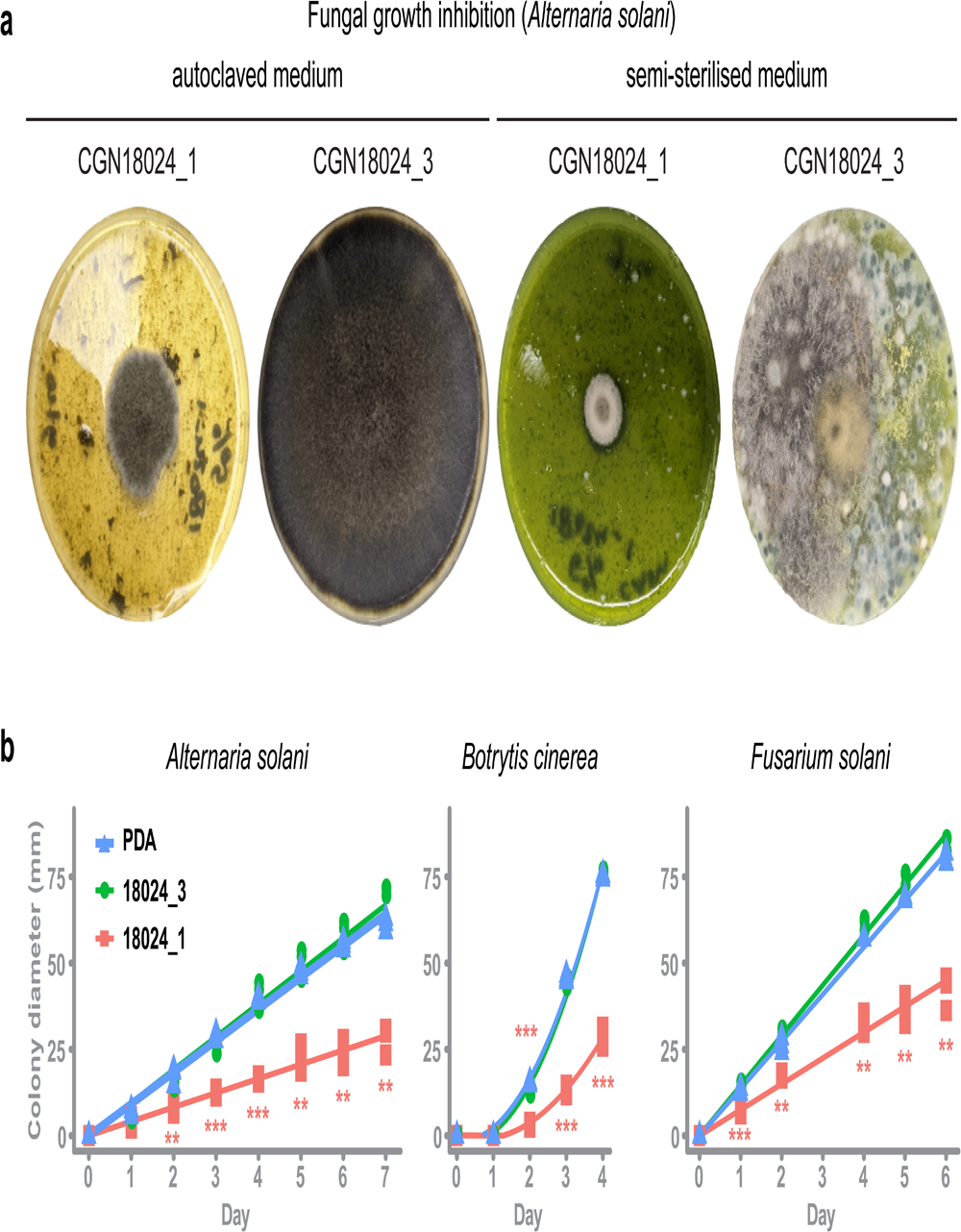
Leaf compounds from resistant S. commersonii inhibit growth of diverse fungi, including pathogens of potato. **A.** Crude leaf extract from CGN18024_1/CGN18024_3 was added to PDA plates (5% w/v) and autoclaved (left) or semi-sterilised for 15 min at 60 °C (right). Growth of *Alternaria solani* altNL03003 was strongly inhibited on PDA plates with autoclaved leaf extract from CGN18024_1 compared to plates with CGN18024_3, as shown on the left two pictures taken at 7 days after placing an agar plug with mycelium of *A. solani* at the centre of each plate. Abundant fungal contamination appeared after 4 days on plates containing semi-sterilised leaf from CGN18024_3, but not on plates containing material from CGN18024_1 (right two pictures). **B.** Growth of potato pathogenic fungi *A. solani* (altNL03003), *B. cinerea* (B05.10) and *F. solani* (1992 vr) was followed by measuring the colony diameter on PDA plates containing autoclaved leaf material from CGN18024_1/CGN18024_3. Growth of all three fungi was measured on PDA plates containing CGN18024_1 (red squares), CGN18024_3 (green circles) or plates with PDA and no leaf material (blue triangles). Significant differences in growth on PDA plates containing plant extract compared to PDA plates without leaf extract are indicated with asterisks (Welch’s Two Sample t-test, \*\**P* < 0.01, \*\*\**P* < 0.001).

To quantify the inhibitory effect of leaves from *S. commersonii* against different fungal pathogens of potato, we performed a growth inhibition assay with *A. solani* (altNL03003 (67)), *Botrytis cinerea* (B05.10 (83)) and *Fusarium solani* (1992 vr). As before, we added 5% (w/v) of leaf material from CGN18024_1 or CGN18024_3 to PDA plates and we placed the fungi at the centre of the plates. We measured colony diameters in the following days and compared it with the growth on PDA plates without leaf extract. Indeed, growth of all three potato pathogens was significantly reduced on medium containing leaf material from CGN18024_1 (Fig 3B), compared to medium containing material from CGN18024_3 or on normal PDA plates. These results indicate that phytoanticipins from the leaves of resistant *S. commersonii* have the potential to protect against diverse fungal pathogens of potato.

### Tetraose steroidal glycoalkaloids from ***S. commersonii*** provide resistance to ***A. solani*** and Colorado potato beetle

*Solanum* leaves usually contain SGAs, which are known phytoanticipins against fungi and other plant parasites (84). *S. tuberosum* typically produces the triose SGAs α-solanine (solanidine-Gal-Glu-Rha) and α-chaconine (solanidine-Glu-Rha-Rha), while five major tetraose SGAs were previously identified in *S. commersonii*: commersonine (demissidine-Gal-Glu-Glu-Glu), dehydrocommersonine (solanidine-Gal-Glu-Glu-Glu), demissine (demissidine-Gal-Glu-Glu-Xyl), dehydrodemissine (solanidine-Gal-Glu-Glu-Xyl) and α-tomatine (tomatidine-Gal-Glu-Glu-Xyl) (84–89). To test if SGAs can explain resistance of *S. commersonii*, we measured SGA content in leaves from Atlantic and susceptible/resistant *S. commersonii* by Ultra High Performance Liquid Chromatography (UPLC) coupled to mass spectrometry (MS). As expected, we could detect the triose SGAs α-solanine and α-chaconine in susceptible *S. tuberosum* cv. Atlantic, but we found a remarkable difference in the SGA profile of resistant and susceptible *S. commersonii*. We detected tetraose SGAs in resistant *S. commersonii* CGN18024_1, whereas susceptible *S. commersonii* CGN18024_3 accumulates triose SGAs (Fig 4A and S3 and S4 Tables). The mass spectra of the four major tetraose SGAs from *S. commersonii* correspond to (dehydro-) commersonine and (dehydro-) demissine, matching the data from previous studies(85, 87–89). Notably, the mass spectra of the two major SGAs from susceptible CGN18024_3 correspond to the triose precursors of these SGAs (solanidine-Gal-Glu-Glu and demissidine-Gal-Glu-Glu respectively) (S3 and S4 Tables). These results suggest that the triose SGAs present in susceptible CGN18024_3 are modified to produce the tetraose SGAs in resistant CGN18024_1, by addition of an extra glucose or xylose moiety.

**Fig 4.**
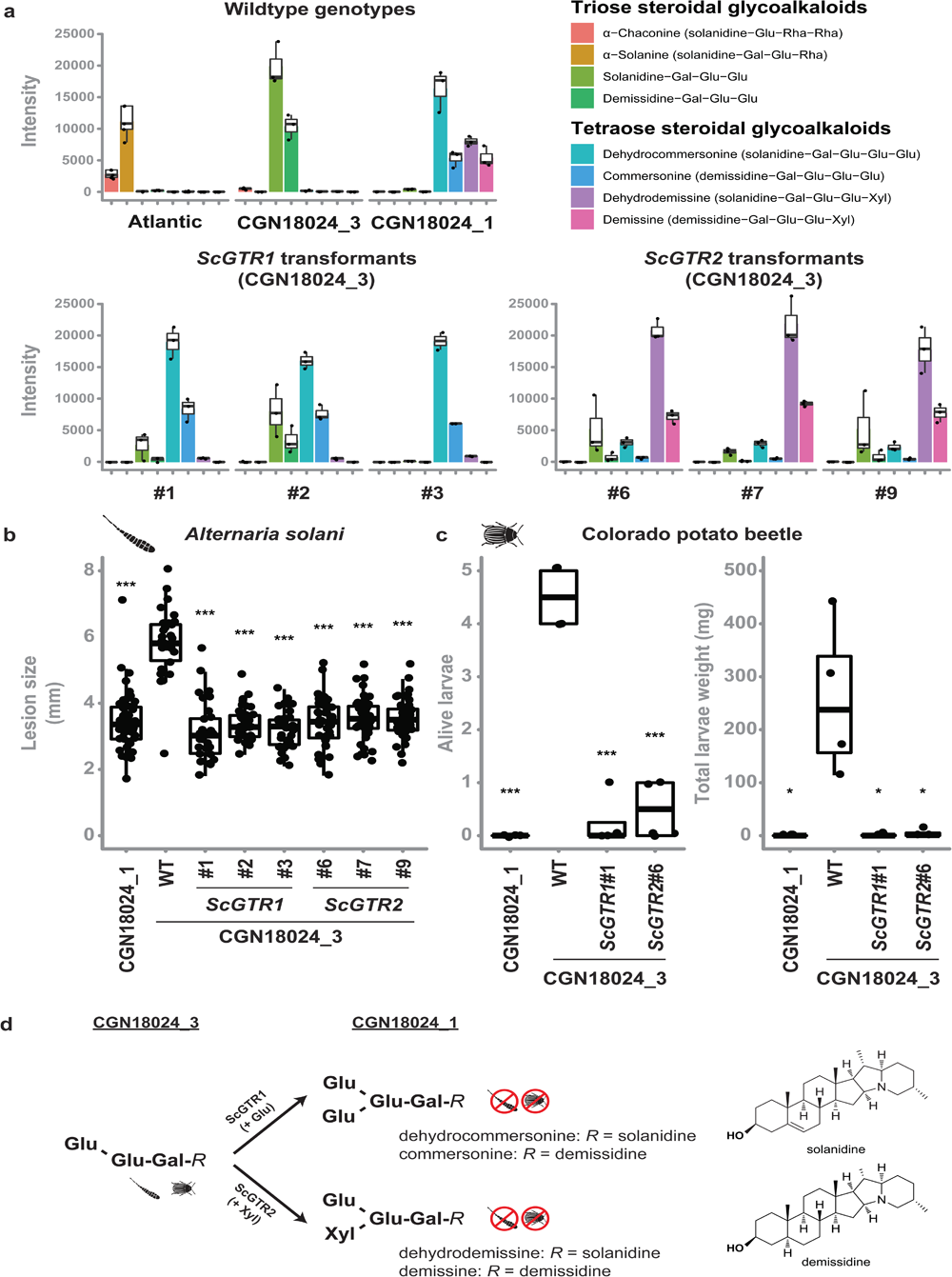
Tetraose steroidal glycoalkaloids from Solanum commersonii provide resistance to Alternaria solani and Colorado potato beetle. Data are visualised with boxplots, with horizonal lines indicating median values and individual measurements plotted on top. **A.** Tetraose steroidal glycoalkaloids (SGAs) were detected in resistant CGN18024_1 and in CGN18024_3 transformed with *ScGTR1*/*ScGTR2*. Susceptible *S. tuberosum* cv. ‘Atlantic’ and wildtype (WT) CGN18024_3 contain only triose SGAs. Overexpression of *ScGTR1* resulted in the addition of a hexose to the triose SGAs from CGN18024_3, resulting in a commertetraose (Gal-Glu-Glu-Glu), while overexpression of *ScGTR2* caused the addition of a pentose, resulting in a lycotetraose (Gal-Glu-Glu-Xyl). **B.** WT CGN18024_1/CGN18024_3 and CGN18024_3 transformants were inoculated with *Alternaria solani* altNL03003. 3 plants of each genotype were tested and 3 leaves per plants were inoculated with 6 10 µl droplets with spore suspension each. Lesions diameters were measured 5 days post inoculation. *ScGTR1* and *ScGTR2* can both complement resistance to *A. solani* in CGN18024_3, as the lesion sizes produced on CGN18024_3 transformants are comparable to resistant CGN18024_1. **C.** 3 plants per genotype were challenged with 5 Colorado potato beetle larvae each. The tetraose SGAs produced by *ScGTR1* and *ScGTR2* can provide resistance to Colorado potato beetle, as indicated by reduced larvae survival and total larvae weight. Significant differences with WT CGN18024_3 are indicated with asterisks (Welch’s Two Sample t-test, \**P* < 0.05, \*\*\**P* < 0.001). **D**. Putative structures of SGAs detected in CGN18024_1 and CGN18024_3, based on previous studies (85, 87–89). CGN18024_3 produces triose SGAs and is susceptible to Colorado potato beetle and *A. solani*. ScGTR1 and ScGTR2 from CGN18024_1 convert these triose SGAs from susceptible *S. commersonii* to tetraose SGAs, through the addition of a glucose or xylose moiety respectively. Both sugar additions can provide resistance to Colorado potato beetle and *A. solani*.

To investigate a possible role for *ScGTR1* and *ScGTR2* in the production of tetraose SGAs from CGN18024_1 and their link to resistance, we generated stable transformants of *ScGTR1* and *ScGTR2* in triose SGA producing CGN18024_3 (S11 Fig). UPLC-MS analysis showed that both *ScGTR1* and *ScGTR2* transformants accumulate tetraose SGAs, while the amount of triose SGAs is markedly reduced (Fig 4A). Strikingly, ScGTR1 and ScGTR2 appear to have different specificities. Overexpression of *ScGTR1* resulted in the addition of a hexose to the triose SGAs from CGN18024_3 (corresponding to a commertetraose), while overexpression of *ScGTR2* caused the addition of a pentose (corresponding to a lycotetraose) (Fig 4A and D). This *in planta* evidence suggests that *ScGTR1* is a glucosyltransferase and that *ScGTR2* is a xylosyltransferase. However, we detect a slight overlap in activity. In addition to the lycotetraose products, we detected small amounts of commertetraose product in *ScGTR1* transformants and vice versa in the *ScGTR2* transformants (Fig 4A and S3 Table). A multivariate Principal Components Analysis (PCA) on the full metabolic profile consisting of all 1,041 detected mass peaks revealed that ScGTR1 and ScGTR2 are highly specific towards SGAs since 75% of the metabolic variation between the transformants and the wild types could be explained by the SGA modifications (S12 Fig). Modifications catalysed by both enzymes can lead to resistance, as *ScGTR1* and *ScGTR2* transformants are both resistant to *A. solani* isolate altNL03003 (Fig 4B). Atlantic *ScGTR1* and *ScGTR2* transformants did not show differences in SGA profile, probably because they contain different triose SGA substrates than found in *S. commersonii* CGN18024_3 (S3 and S4 Tables).

Leptine and dehydrocommersonine SGAs from wild potato relatives have previously been linked to resistance to insects such as CPB (44, 47–51, 90). To see if the SGAs from *S. commersonii* can protect against insects as well, we performed a test with larvae of a CPB genotype collected in the Netherlands on wildtype CGN18024_1/CGN18024_3 and on CGN18024_3 transformed with *ScGTR1* or *ScGTR2* (Fig 4B). Wildtype CGN18024_3 is susceptible to the CPB genotype that was tested, but CGN18024_1 and CGN18024_3 transformed with *ScGTR1* or *ScGTR2* are resistant, as illustrated by a very low larvae weight and survival (Fig 4C). Thus, the conversion of triose SGAs from CGN18024_3 to tetraose SGAs produced by CGN18024_1, carried out by both ScGTR1 and ScGTR2, can provide protection against *A. solani* as well as CPB (Fig 4A-D).

## Discussion

In this study, we set out to characterise resistance of *S. commersonii* to *A. solani*. We showed that it is caused by a single dominant locus containing two GT candidate resistance genes. Both GTs are involved in the production of tetraose SGAs in *S. commersonii*, but they transfer distinct sugars. Both modifications can cause resistance to *A. solani*. We provide *in vitro* evidence to show that the tetraose SGAs from *S. commersonii* have the potential to protect against other fungi besides *A. solani* and we demonstrate that plants producing the compounds are resistant to CPB. Collectively, our data link the tetraose SGAs from *S. commersonii* to disease and pest resistance.

It is known that specialised metabolites from plants can act in plant defence and compounds with antimicrobial effects have been characterised in many different plant species (26, 27, 91). They may also influence other aspects of the crop, such as flavour or taste and they can have dietary benefits or be toxic to humans. SGAs from potato can cause risks for human health, but a total SGA content of less than 200 mg/kg is generally considered to be safe for human consumption (84, 92–94). Potato breeders generally try to reduce SGA content in tubers, to prevent problems with toxicity and to meet safety regulations, but they do not usually consider the effect on disease resistance. There is not much known about how modifications to SGAs of potato affect human toxicity and resistance to biotic stress, but additional knowledge on this topic could help breeders to optimize the metabolite profile of their cultivars (95).

Biosynthesis of SGAs in *Solanum* is controlled by many genes. The discovery of *S. commersonii* genotypes with and without tetraose SGAs provides us with unique insight in the role of these compounds in plant immunity. Similar compounds are produced in *Solanum* species such as *S. chacoense*, *S. chomatophilum*, *S. oplocense*, *S. paucisectum* and *S. piurae*, which may explain why these (or their descendants) display resistance to *A. solani* or CPB (48–50, 96, 97). The compounds that are found in resistant *S. commersonii* are an interesting combination of a solanidine or demissidine aglycone and a lycotetraose or commertetraose sugar moiety. Solanidine forms the aglycone backbone of α-solanine and α-chaconine from potato, while the lycotetraose decoration is found on α-tomatine from tomato (87, 98). The biosynthesis pathways leading to the production of these major SGAs from cultivated potato and tomato have largely been elucidated in recent years and it was found that the underlying genes occur in conserved clusters (76, 98). This knowledge and the similarities between SGAs from *S. commersonii* and cultivated potato/tomato will help to identify the missing genes from the pathway through comparative genomics.

The broad-spectrum activity of tetraose SGAs is attractive, but this non-specificity also presents a risk. The antifungal and anti-insect activity of SGAs from *S. commersonii* is not restricted to potato pathogens and pests, but could also affect beneficial or commensal micro-organisms or other animals that feed on plants (43, 99). In tomato fruit, α-tomatine is converted to esculeoside A during fruit ripening in a natural detoxification process from the plant (100, 101) to facilitate dispersal of the seeds by foraging animals. Unintended toxic effects of SGAs should also be taken into account when used in resistance breeding.

Studies on α-tomatine and avenacin A-1 show that changes to the sugar moiety of these saponins from tomato and oat respectively, can affect their toxicity (46, 102–104). Tomato and oat pathogens produce enzymes that can detoxify these compounds through removal of one or more glycosyl groups (46, 52, 53, 105–107). The degradation products of saponins can also suppress plant defence responses (108, 109). Conversely, here we show that the resistance of *S. commersonii* is based on the addition of a glycosyl group to a triose saponin from *S. commersonii.* There is large variation in both the aglycone and the sugar moiety of SGAs from wild *Solanum*, with likely over 100 distinct SGAs produced in tubers (87) (110). This diversity suggests a pressure to evolve novel molecules, possibly to resist detoxification or other tolerance mechanisms, reminiscent of the molecular arms race that drives the evolution of plant immune receptors (14). Thus, wild *Solanum* germplasm is not only a rich source of immune receptors, it also provides a promising source of natural defence molecules. Studies of how pathogens that naturally occur on *S. commersonii*, or other *Solanum* species producing tetraose SGAs, can tolerate SGAs produced by their hosts could help judge the durability of this type of resistance.

As crops are usually affected by multiple diseases and pests, significant reduction of pesticide use can only be achieved if plants are naturally protected against a range of pathogen species and pests. Different strategies towards this goal have been proposed and our study underlines the potential of defence compounds that are naturally produced by plants. The fact that genes for specialised plant metabolites can occur in biosynthetic gene clusters (76, 111–113), means that introgression breeding could help to move these compounds from wild relatives to crop species. We had already created *S. commersonii* x *S. tuberosum* hybrids with resistance to early blight in a previous study (58), but it is clear that potential negative effects of SGA variants on human health and the environment should be considered before these can be developed into a cultivar.

Additional insight in the biosynthesis pathway of the tetraose SGAs produced by *S. commersonii* would make it possible to employ them through metabolic engineering and allow for a more precise control of the amounts that are produced and in which tissues (91). Alternatively, the defence compounds could be produced in non-crop plants or other organisms and applied on crops as biological protectants. Studies on how natural defence compounds are produced in different plant tissues, their toxicity and how they are detoxified, combined with studies on how different modifications ultimately affect plant immunity and toxicity, are essential to employ them in a safe and effective manner. Such studies at the interface of plant immunity and metabolism can help to design innovative solutions to complement existing resistance breeding strategies and improve sustainability of our food production.

## Methods

A brief method description is given below, full details on methods can be found in S2 File. The primers used in this study are listed in S5-7 Tables.

### Genome assembly and separation of haplotypes covering resistance region

ONT reads were filtered using Filtlong v0.2.0 (https://github.com/rrwick/Filtlong) with --min_length 1000 and --keep_percent 90. Adapter sequences were removed using Porechop (114). Fastq files were converted to Fasta using seqtk v1.3 (https://github.com/lh3/seqtk). Assembly was performed with smartdenovo (https://github.com/ruanjue/smartdenovo/) and a k-mer size of 17, with the option for generating a consensus sequence enabled. ONT reads were mapped back to the assembly using minimap2 v2.17 (115) and used for polishing with racon v1.4.3 (116) using default settings. DNBseq reads were mapped to the resulting sequence using bwa mem v0.7.17 (117) and used for a second round of polishing with racon v1.4.3. This procedure to polish the assembly using DNBseq reads was repeated once. ONT reads were mapped back to the polished CGN18024_1 assembly using minimap2 v2.17 (115). The alignment was inspected using IGV v2.6.3 (118) to identify polymorphisms for new markers and marker information was used to identify ONT reads representative for both haplotypes spanning the resistance region of CGN18024_1. Bedtools v2.25.0 (119) was used extract the resistance region from the reads and to mask the corresponding region from the original CGN18024_1 assembly. The extracted resistance regions from both reads were appended to the assembly and the polishing procedure described above was repeated to prepare a polished genome assembly of CGN18024_1, containing a sequence of both haplotypes covering the resistance region. Quality of the genome was assessed using quast v5.0.2 with --eukaryote --large (120).

### Transient disease assay

Agroinfiltration was performed as described previously using *Agrobacterium tumefaciens* strain AGL1 (82, 121). Agrobacterium suspensions were used at an OD600 of 0.3 to infiltrate fully expanded leaves of 3-week-old CGN18024_1, CGN18024_3 and *S. tuberosum* cv. Atlantic. *ScGTR1*, *ScGTR2*, *ScGTS* and pK7WG2-empty were combined as four separate spots on the same leaf and the infiltrated areas were encircled with permanent marker. The plants were transferred to a climate cell 48 h after agroinfiltration and each infiltrated area was inoculated with *A. solani* by pipetting a 10 µl droplet of spore suspension (1 x 10 conidia/ml) at the centre of each spot. Lesion diameters were measured 5 days post inoculation. Eight plants were tested of each genotype, using three leaves per plant.

### Fungal growth inhibition assays

Mature leaf material from 5-week-old plants was extracted in phosphate-buffered saline (PBS) buffer using a T25 Ultra Turrax disperser (IKA) and supplemented to obtain a 5% w/v suspension in PDA and autoclaved (20 min at 121C), or added to PDA after autoclaving, followed by an incubation step for 15 min at 60C to semi-sterilise the medium. The medium was poured into Petri dishes. Small agar plugs containing mycelium from *A. solani* (altNL03003) or *F. solani* (1992 vr) were placed at the centre of each plate and the plates were incubated at 25C in the dark. Similarly, approximately 100 spores of *B. cinerea* B05.10 (83) were pipetted at the centre of PDA plates containing the different leaf extracts and the plates were incubated at room temperature in the dark. 3 plates per fungal isolate/leaf extract combination were prepared and colony diameters were measured daily using a digital calliper.

### Data analysis

Data were analysed in RStudio (R version 4.02) (122, 123), using the tidyverse package (124). Most figures were generated using ggplot2 (125), but genomic data were visualised using Gviz and Bioconductor (126). PCA was performed using PAST3 software (https://past.en.lo4d.com/windows). *P* values for comparisons between means of different groups were calculated in R using Welch’s Two Sample t-test.

### Data availability

RNAseq data from the BSR-Seq experiment was deposited in the NCBI Sequence Read Archive with BioProject ID PRJNA792513 (Sequencing Read Archive accession IDs SRR17334110, SRR17334111, SRR17334112 and SRR17334113). Raw reads used in the assembly of the CGN18024_1 genome were deposited with BioProject ID PPRJNA789120 (Sequencing Read Archive accession IDs SRR17348659 and SRR17348660). The assembled genome sequence of CGN18024_1 was archived in GenBank under accession number JAJTWQ000000000. Sequences of *ScGTR1* and *ScGTR2* were deposited in GenBank under accession numbers OM830430 and OM830431. Numerical data underlying the figures of this manuscript are included in S1 File.

## Acknowledgements

This research was funded by the J.R. Simplot Company, we especially thank Craig Richael for his support and useful discussions. We thank Dirk Jan Huigen and Henk Meurs for taking care of the plants in the greenhouse and Jack Vossen for providing us with *F. solani* isolate 1992 vr from the collection of Biointeractions and Plant Health (Theo van der Lee, WUR). Jan van Kan and Yaohua You for insightful discussions and for providing us with *B. cinerea* isolate B05.10. Evert Jacobsen for his feedback on the manuscript. Martijn van Kaauwen and Richard Finkers for bioinformatics support. P.J.W. thanks Andrea Lorena Herrera for her support and helpful talks about specialised metabolites.

## Supporting information

**S1 Fig. Resistance of *S commersonii* genotypes CGN18024_1 and CGN18024_3 against different isolates of *A. solani*.** CGN18024_1 and CGN18024_3 were inoculated with A. solani altNL03003 (the reference isolate used throughout the manuscript), altNL21001 (isolated from a potato field in the Netherlands in 2021) and ConR1H (harvested from a potato field in Idaho, USA in 2015). CGN18024_1 is resistant against all three isolates, whereas CGN18024_3 is susceptible

**S2 Fig. Resistance from *S. commersonii* to *A. solani* is mapped to the top of chromosome** F**1**il**2**te. red SNPs from bulked segregant RNAseq analysis (BSA-RNAseq) are plotted in 100 kb windows on chromosome 12 of the DMv4.03 genome at the top of the figure. A selection of SNPs (‘A1’-‘A10’ and ‘B1’-‘B4’) was used as markers in high resolution melting (HRM) analysis to genotype resistant S commersonii parent CGN18024_1 and susceptible parent CGN18024_3 from the AJW12 mapping population as well as progeny used in BSA-RNAseq. HRM analysis led to the identification of recombinants AJW12_13, AJW12_18, AJW12_23 and AJW12_29. Recombinant AJW12_13 (susceptible to ***A.*** solani) and recombinant AJW12_29 (resistant to A. solani) are used to map the resistance locus from S. commersonii to a window of approximately 3 Mb at the top of chromosome 12, delimited by marker ‘B3’.

**S3 Fig. Overview of marker 817K.** Integrated Genomics Viewer (IGV) snapshot of Oxford Nanopore Technology (ONT) reads aligned to the genome of S. commersonii CGN18024_1. An Insertion/Deletion (InDel) of 254 bp is observed at approximately 817 kb of contig utg1998 that covers the resistance region. Primers were designed flanking the InDel to develop marker 817K.

**S4 Fig. Overview of marker 807K.** Integrated Genomics Viewer (IGV) snapshot of Oxford Nanopore Technology (ONT) reads aligned to the genome of S. commersonii CGN18024_1. An Insertion/Deletion (InDel) of 310 bp is observed at approximately 807 kb of contig utg1998 that covers the resistance region. Primers were designed flanking the InDel to develop marker 807K.

**S5 Fig. Overview of marker 797K.** Integrated Genomics Viewer (IGV) snapshot of Oxford Nanopore Technology (ONT) reads aligned to the genome of S. commersonii CGN18024_1. An Insertion/Deletion (InDel) of 6 bp is observed at approximately 797 kb of contig utg1998 that covers the resistance region. Primers were designed flanking the InDel to develop marker 797K.

**S6 Fig. Overview of marker 764K.** Integrated Genomics Viewer (IGV) snapshot of Oxford Nanopore Technology (ONT) reads aligned to the genome of S. commersonii CGN18024_1. An Insertion/Deletion (InDel) of 47 bp is observed at approximately 764 kb of contig utg1998 that covers the resistance region. Primers were designed flanking the InDel to develop marker 764K.

**S7 F ig. Fine m apping t h e resistance l ocus in CGN18024_1.** he. Solyntus and CGN18024_1 genome were used to screen for recombinants among progeny from a cross between resistant CGN18024_1 and susceptible CGN18024_3. Physical locations of the markers on the DMv4.04, Solyntus and CGN18024_1 genome are indicated at the top of the figure. Recombinants that were identified were tested for resistance to A. solani to fine map the resistance region. Recombinants 2-G10 (resistant, R), 14-F06 and 14-C12 (both susceptible, S) are used to delimit the resistance region between markers 817K and 797K, corresponding to a region of 20 kb in the CGN18024_1 genome.

**S8 Fig. Early blight disease symptoms on key recombinants.** The picture shows lesions of representative leaves of key recombinants at 5 days post drop-inoculation with spores of A. solani.

**S9 Fig. Alignment of putative *S. commersonii* glycosyltransferases (ScGTs) linked to resistance.** ScGTR1, ScGTR2 and ScGTS show high similarity, but the GT encoded by the susceptible haplotype (ScGTS) contains a mutation that leads to a truncated protein.

**S10 Fig. Comparative phylogenetic analysis of glycosyltransferases with a known function (S2 Table).** The phylogenetic tree is constructed using the maximum likelihood method (100 bootstraps). ScGTR1 and ScGTR2 are indicated with arrows and GTs with a previously characterised role in SGA biosynthesis are marked with asterisks. Direct homologs of these SGA GTs (based on identity and synteny) derived from the CGN18024_1 genome are included in the analysis (names starting with ‘SCM’).

**S11 Fig. Validation of *ScGTR1* and *ScGTR2* transformants using PCR.** Gel electrophoresis of PCR amplicons produced by primer combinations p35S + ScGTR1sr3 (ScGTR1), p35S + ScGTR2sr3 (ScGTR2) and ef1αF1 + ef1αR1 (EF1α) using genomic DNA template of wildtype CGN18024_3 (WT) and ScGTR1/ScGTR2 CGN18024_3 transformants.

**S12 Fig. Principal Component Analysis (PCA) on *Solanum commersonii* genotypes and transformants.** PCA based on 1041 mass peaks detected by UPLC-MS in leaves of ScGTR1 (red dots) and ScGTR2 (red squares) transformants compared to the corresponding susceptible wildtype Solanum commersonii CGN18024_3 (blue circles) and resistant CGN18024_1 (yellow circles). 75% of the total metabolic variation between the groups is explained by the 1st and the 2nd PC, mostly loaded by variation between tri-and tetraglycosylated steroidal glycoalkaloids. S/DhS - Solanidine/demissidine.

**S1 Table. *Solanum commersonii* and *Solanum malmeanum* accessions used in this study.** Accessions were obtained from the Centre for Genetic Resources, the Netherlands (CGN WUR). 2-3 genotypes from each accession were used in the disease screen with A. solani.

**S2 Table. Overview of characterised glycosyltransferases used in comparative phylogenetic analysis (S9 Fig).** Glycosyltransferases (GTs) with a known function are taken from Bowles et al. (2005), McCue et al. (2005, 2006, 2007), Masada et al. (2009), Itkin et al. (2011, 2013) and Tikunov et al. (2013) (71–78)

**S3 Table. Putative identities and relative contents of SGAs in different potato genotypes.** intensities (3 replicates per genotype) are presented as a percentage of the maximum signal intensity.

**S4 Table. Overview of the steroidal glycoalkaloids detected in our study.** RT - retention time, [M-H+FA]- - mass of a molecular ion at negative ionization mode (all alkaloids were represented by formic acid adduct ions); [M+H]+ - mass of a molecular ion at negative ionization mode; Putative structure - putative combination of aglycones and sugar moieties deduced by comparing the fragmentation spectrum derived at positive ionization with previous studies (85, 87–89); Fragmentation spectra derived using positive ionization: P - parent ion or P-fragment(s) loss.

**S5 Table.** Overview of primers used to map the resistance region.

**S6 Table.** Overview of primers used to clone candidate resistance genes.

**S7 Table.** Overview of primers used to validate transformants.

**S1 File**. Numerical data underlying the figures of this manuscript.

**S2 File**. Full information on methods.

