## Supplementary figures and images for "Tetraose steroidal glycoalkaloids from potato provide resistance against *Alternaria solani* and Colorado potato beetle"

### S1Fig

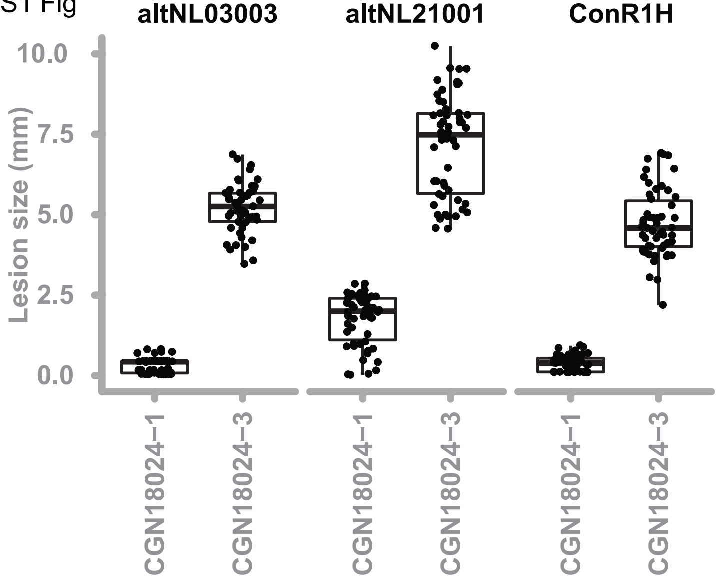

### S2Fig

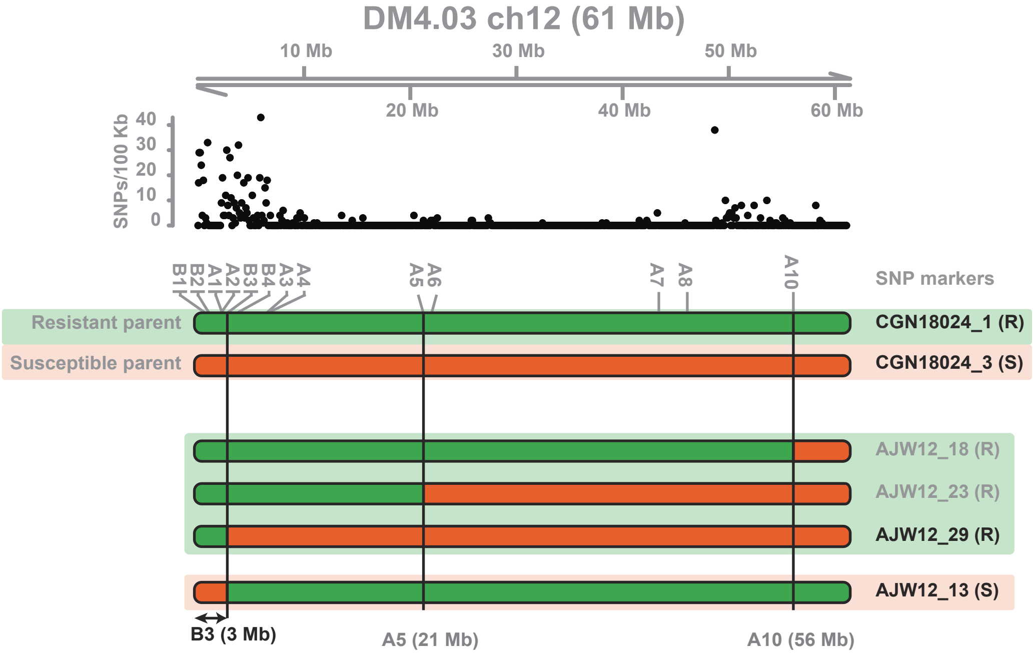

### S7Fig

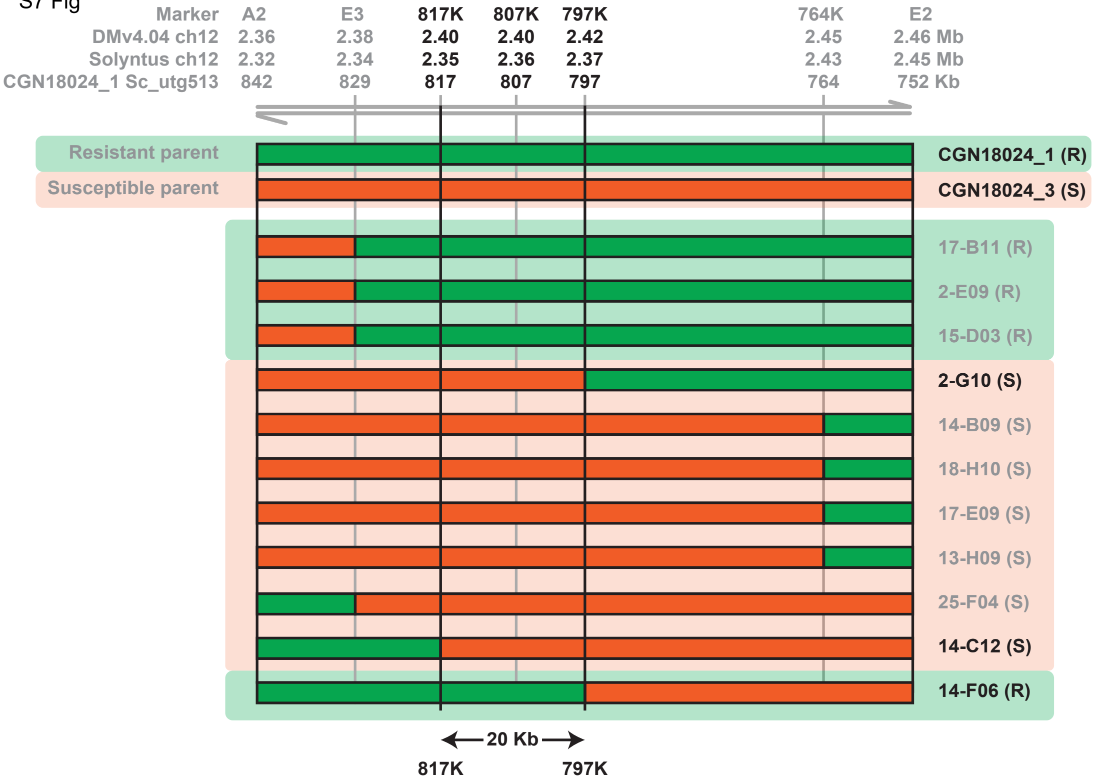

### S8Fig

S8 Fig

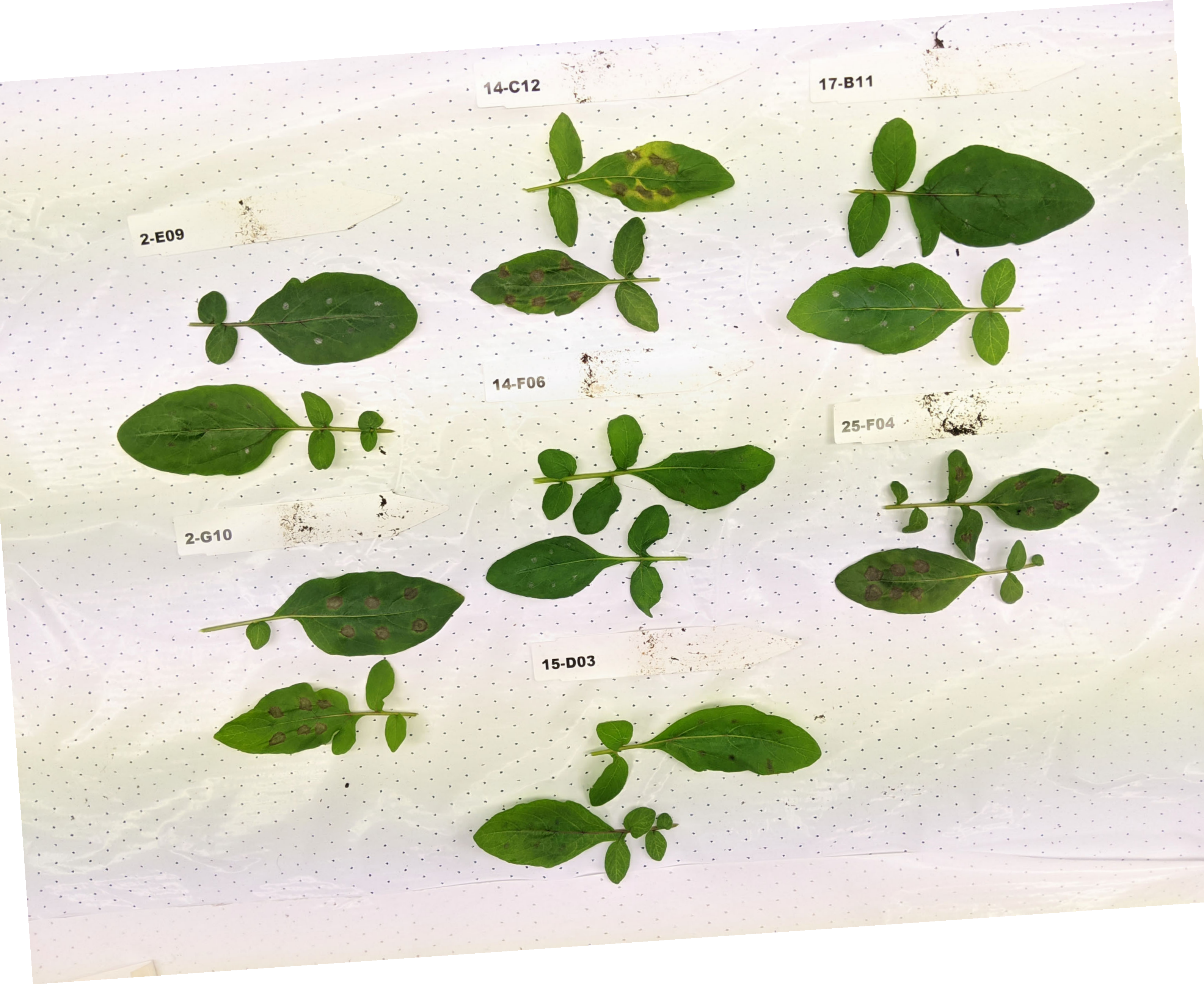

### S10Fig

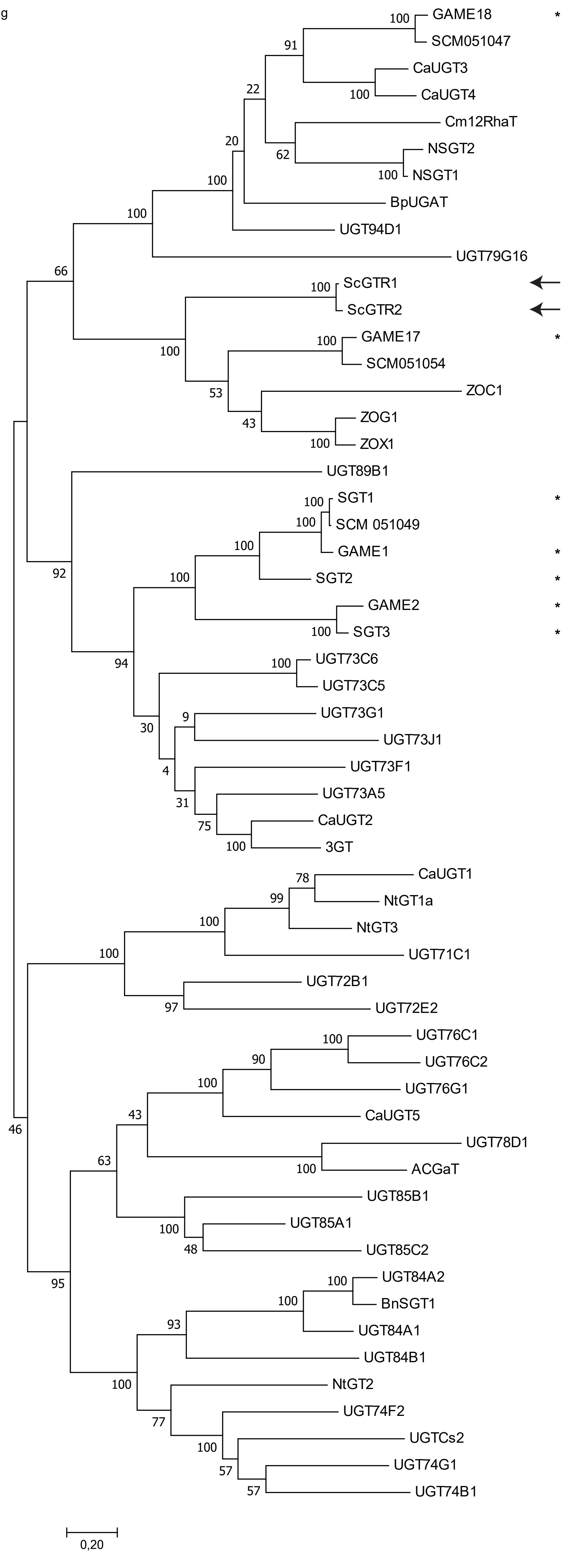

### S11Fig

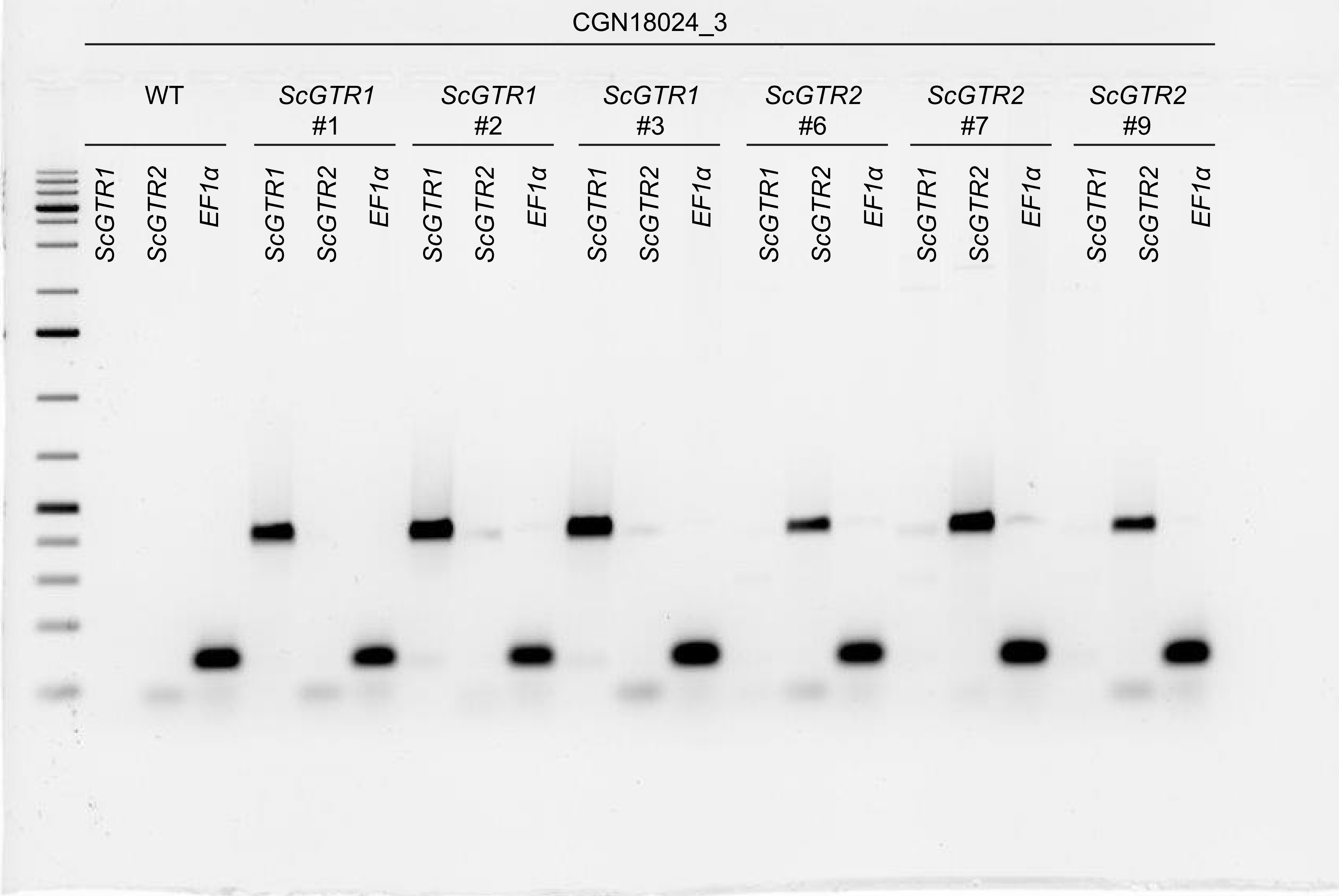

### S12Fig

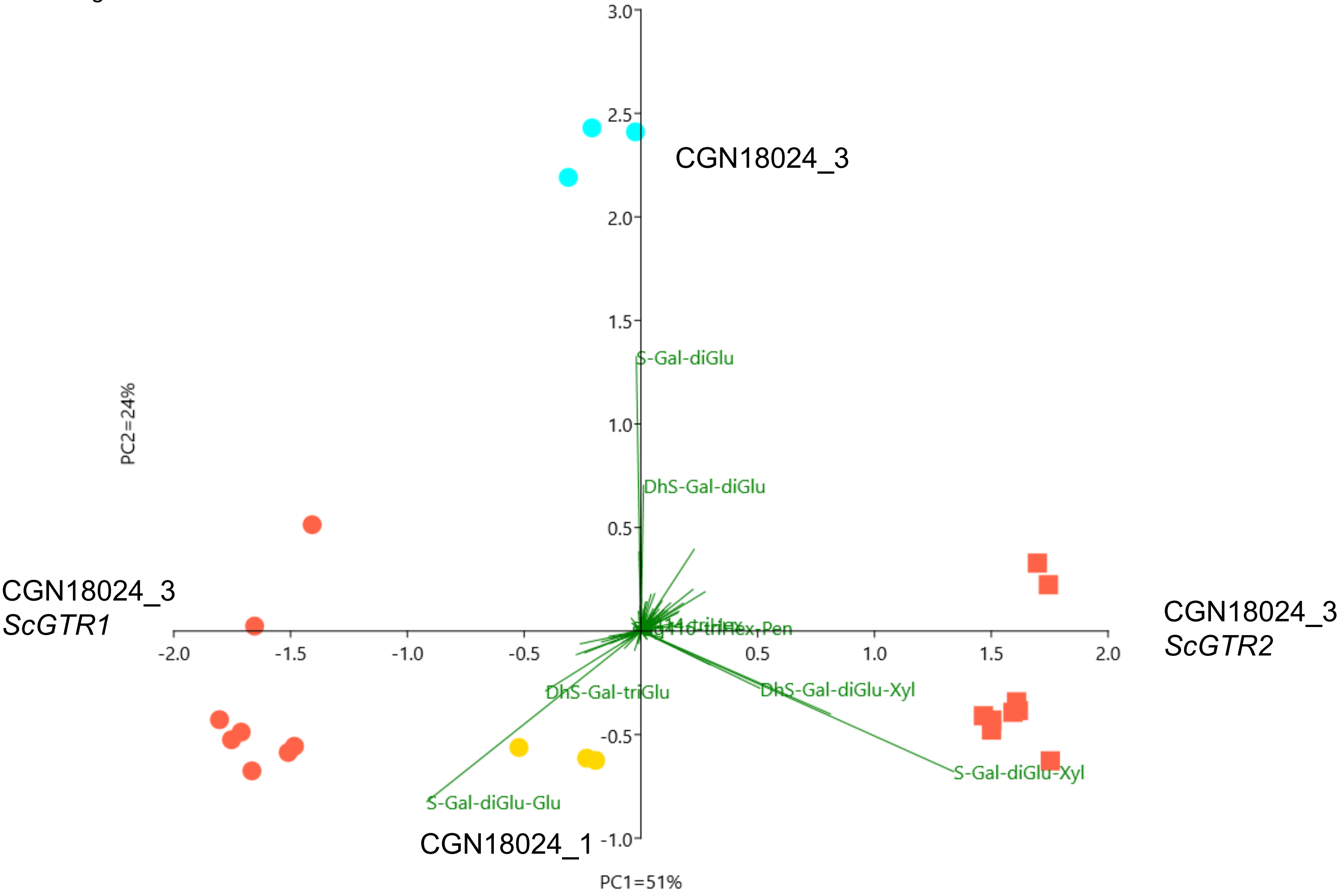
