## Supplementary material for "Tetraose steroidal glycoalkaloids from potato provide resistance against *Alternaria solani* and Colorado potato beetle": S2File

Supplementary methods

**Plant material.** Seeds from *S. commersonii* and *S. malmeanum accessions* (S1 Table) were obtained from the CGN germplasm collection (Wageningen, the Netherlands). Seeds were sterilised by washing them in 70% ethanol, followed by a 15 min incubation in a 1.2% sodium hypochlorite solution. Sterilised seeds were rinsed 3 times in sterile tap water and sown out on MS20 medium and incubated in the dark until germinated.

The mapping population was generated by crossing *S. commersonii* CGN18024_1 with CGN18024_3 and vice versa. Ripe berries were harvested about 6 weeks after pollination. Seeds were harvested from the ripe berries, washed with tap water and dried at room temperature on filter paper for 2 weeks. Dry seeds were stored at 4 °C until use.

All plants were maintained in tissue culture on MS20 medium (4.4 g Murashige and Skoog basal salt mixture including vitamins, 20 g sucrose and 8 g/L micro agar, pH=5.8)[1]. Fresh shoots were propagated two weeks prior to transferring plants to soil. Plants were grown in a greenhouse under long-day conditions (16 h light/8 h dark).

**Production of inoculum from *A. solani*.** Spores from *A. solani* isolate CBS 143772/altNL03003 were produced as described in Wolters *et al.* (2019). The isolate was maintained on potato dextrose agar (1). Small plugs containing mycelium of *A. solani* were transferred to autoclaved V8 medium (5x diluted V8 juice) and incubated in a shaking incubator at 25 °C for five days. The resulting culture was poured into PDA petri dishes and incubated without lid in an incubator equipped with blacklight fluorescent tubes (Philips TL-D 18 W BLB) at 25 °C (12 h dark/12 h blacklight) for 3 days. The petri dishes containing spores were covered and kept in the dark at 25 °C for about a week. Inoculum was prepared by pouring tap water into the petri dishes and gently scraping conidia from the plates using a plate spreader. The conidial suspension was filtered through a tea strainer and collected in a 50 ml tube. Spore concentration was calculated using a haemocytometer. Conidia were left to settle to the bottom of the 50 ml tube and resuspended in ½ strength potato dextrose broth supplemented with 0.3% micro-agar to obtain a final suspension of 1 × 10^5^ conidia/ml for drop-inoculation, or resuspended in ½ strength PDB for spray-inoculation.

**Early blight testing.** 5-week-old plants were transferred to a climate cell two days before the disease assay (25 °C, RH=70% and 16 h light/8 h dark). The plants were placed inside a transparent plastic tent that also contained an atomiser (Condair 505). 3 replicates per genotype were included and genotypes were equally divided over 3 blocks to maintain comparable distances to the atomiser. Three compound leaves per plant were inoculated with six 10 µl droplets of spore suspension each, or complete plants were spray inoculated (1 × 10^5^ conidia/ml). Lights were switched off and the atomiser was turned on after inoculation. The normal light regime was resumed the next morning and the atomiser was only switched on during subsequent night. Lesion diameters were measured 5 days post inoculation using a digital calliper.

**Isolation of nucleic acids and sequencing.** RNA isolations were performed from leaf material that was harvested from fully expanded leaves of 3-week-old CGN18024_1 and CGN18024_3 and from young leaves from the top of the plant of 3-week-old progeny derived from the cross between these genotypes and of transformants. RNA was extracted using the RNeasy Plant Mini Kit following manufacturer's instructions, including an on-column DNase treatment (Qiagen). RNA sequencing was performed on the Illumina platform (PE150) by Novogene (United Kingdom), using around 4 µg of RNA.

Genomic DNA was isolated using the DNeasy Plant Mini Kit (Qiagen) or in a 96-wells format (2). High molecular weight DNA was isolated from young leaves of CGN18024_1 and CGN18024_3 as described previously (3, 4). Quality and integrity of RNA and DNA samples was assessed using nanodrop, Qubit and gel electrophoresis. ONT sequencing was performed on a Nanopore GridION system, using 3 flow cells, using about 1 ug of DNA per flow cell and a run-time of 72 hr. Approximately 4 ug of genomic DNA was sent to BGI Europe for sequencing on the DNBseq platform.

**BSA-RNAseq analysis.** BSA-RNAseq reads were cleaned with fastp (5) and aligned to the CGN18024_1, DMv4.03 and Solyntus genomes with STAR v2.7.3a (6), using default settings. The alignment files were sorted and indexed using SAMtools v1.9 (7). SAMtools mpileup was used in combination with VarScan v2.3.9 mpileup2SNP (8) with –strandfilter 0 and --output-vcf 1, for SNP calling and to prepare Variant Call Format (VCF) files. To identify putative SNPs linked to resistance, a previously described script was used (9). SNPs that were heterozygous in resistant parent and progeny (SNP frequency of 40-60%), but homozygous or absent in the susceptible parent and progeny (0-10% or 90-100%), were retained (File S1) and used to develop HRM markers.

**Development of markers and genotyping.** Bedtools v2.25.0 (10) was used to extract the regions surrounding polymorphisms from the DMv4.03, Solyntus and CGN18024_1 genomes. Primers were designed using BatchPrimer3 (11) (S5 Table). HRM markers were amplified with Phire Hot Start II DNA Polymerase (Thermo Fisher Scientific) and genotyped on a LightScanner System (Bio Fire) as described previously (2). InDel markers were amplified using DreamTaq DNA Polymerase (Thermo Fisher Scientific) and visualised using gel electrophoresis following standard laboratory protocols.

**Comparing haplotypes covering resistance region.** Genes were predicted using the funannotate v1.7.4 (<https://github.com/nextgenusfs/funannotate/>) pipeline. Briefly, funannotate was used to sort and mask the genome and training was performed using the BSA-RNAseq data (--max_intronlen 10000). Gene prediction was prepared using the --optimize_augustus --organism other and --max_intronlen 10000 options. The two haplotypes covering the resistance region were compared using nucmer and visualised using mummerplot from the MUMmer4 package (12).

**Cloning.** *ScGTR1*, *ScGTR2* and *ScGTS* were amplified from genomic DNA from CGN18024_1 using Phusion Polymerase (New England BioLabs) and the primers listed in S6 Table following standard laboratory protocols. cacc was included at the 5’ end of each forward primer to facilitate cloning in the pENTR D-TOPO vector (Thermo Fisher Scientific), following manufacturer's instructions. Insert sequences were validated through Sanger sequencing (Macrogen Europe). The genes were cloned into the pK7WG2 vector (13) using Gateway LR Clonase II (Thermo Fisher Scientific) following manufacturer's instructions and transformed to electrocompetent *A. tumefaciens* AGL1 (14) containing the helper plasmid pVirG (15).

**Potato transformation.** Internodes from *in vitro* grown plants were used to generate stable transformants using previously described methods (16, 17). Transformants were selected on MS20 containing 100 ug/ml kanamycin. Successful transformants were characterised using primers listed in S7 Table.

**SGA measurements.** Mature leaves were harvested from 3 different 5-week-old plants of each genotype in 2 ml tubes containing 2 steel beads and flash frozen in liquid nitrogen. Leaf material was ground using a TissueLyser II bead mill (Qiagen). Approximately 100 mg of ground leaf material was extracted in 1 ml of 70% methanol and 0.1% formic acid (FA). Samples were vortexed and sonicated for 15 min in an Ultrasonic Cleaner (VWR). Samples were vortexed once more and centrifuged for 15 min in a tabletop centrifuge at 17,000 g. The supernatant was passed through 0.45 µm syringe filters (BGB) and diluted 5x. The extracts were separated on Acquity UPLC HSS T3 1.8 um (2.1x150 mm) column Acquity UPLC H Class Plus system (Waters). The separation was performed using the following water + 0.1 formic acid/acetonitrile + 0.1% formic acid gradient: initial – A/B = 95/5%, 65/35% - 14 min, 55,45% - 20 min, 15/85% - 24 min, 95/5% - 25 min, 95/5% - 30 min. The MS data were acquired using an Acquity QDa mass spectrometer (Waters) in negative and positive mode (in separate runs) from 150 – 1250 Da, cone voltage 15V, capillary voltage 0.8kV at 2 scans/second acquisition rate. The raw chromatograms were subjected to full spectra alignment using Metalign software (<https://www.wur.nl/en/show/MetAlign-1.htm>). SGAs were putatively identified using MS fragmentation patterns (S3 and S4 Tables), which were compared with MS information available in literature (18-21).

**Colorado potato beetle test.** CPB was reared on *S. tuberosum* cultivar ‘Bintje’ in insect rearing cages in a greenhouse compartment at 25/23°C and 16/8 h light/dark photoperiod and 70% relative humidity. Freshly laid egg packages were removed from the rearing every day, and once hatched, one-day old larvae were used for the experiment. Resistance to CBP was measured by assessing mortality and weight of CPB larvae on 3 plants of each genotype in a non-choice assay. At the start of the experiment, five one-day-old larvae were placed in a clip-cage on a leaf and an insect sleeve was used to enclose every plant to restrict the larvae to the plant. Larvae were able to feed for nine days, after which surviving larvae were counted and weighed on a scale. If less than five larvae were found on the plant, the remainder was assumed dead.
