## Supplementary material for "Tetraose steroidal glycoalkaloids from potato provide resistance against *Alternaria solani* and Colorado potato beetle": S3-6Fig

S3 Fig

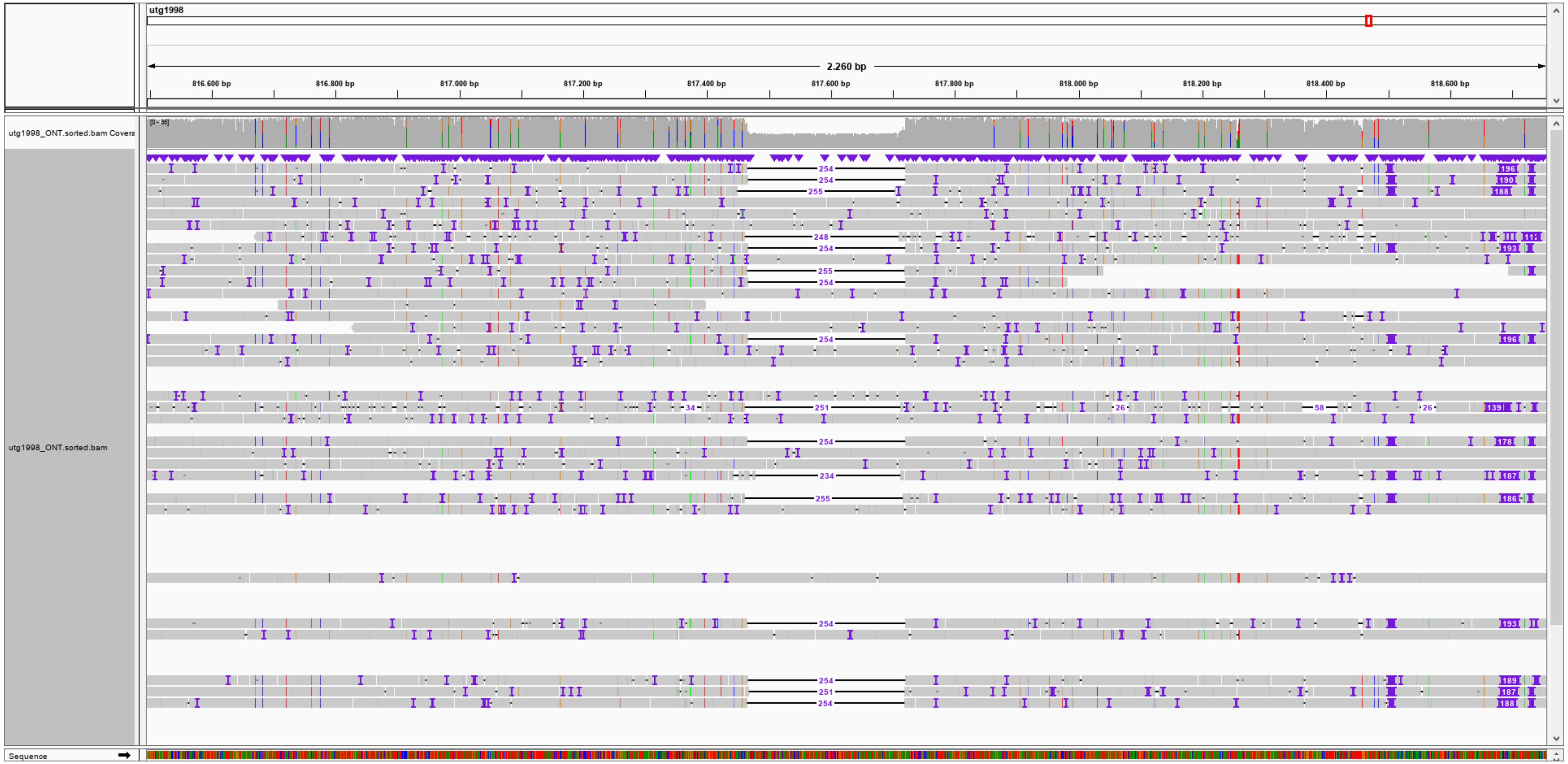


**Overview of marker 817K**

S4 Fig

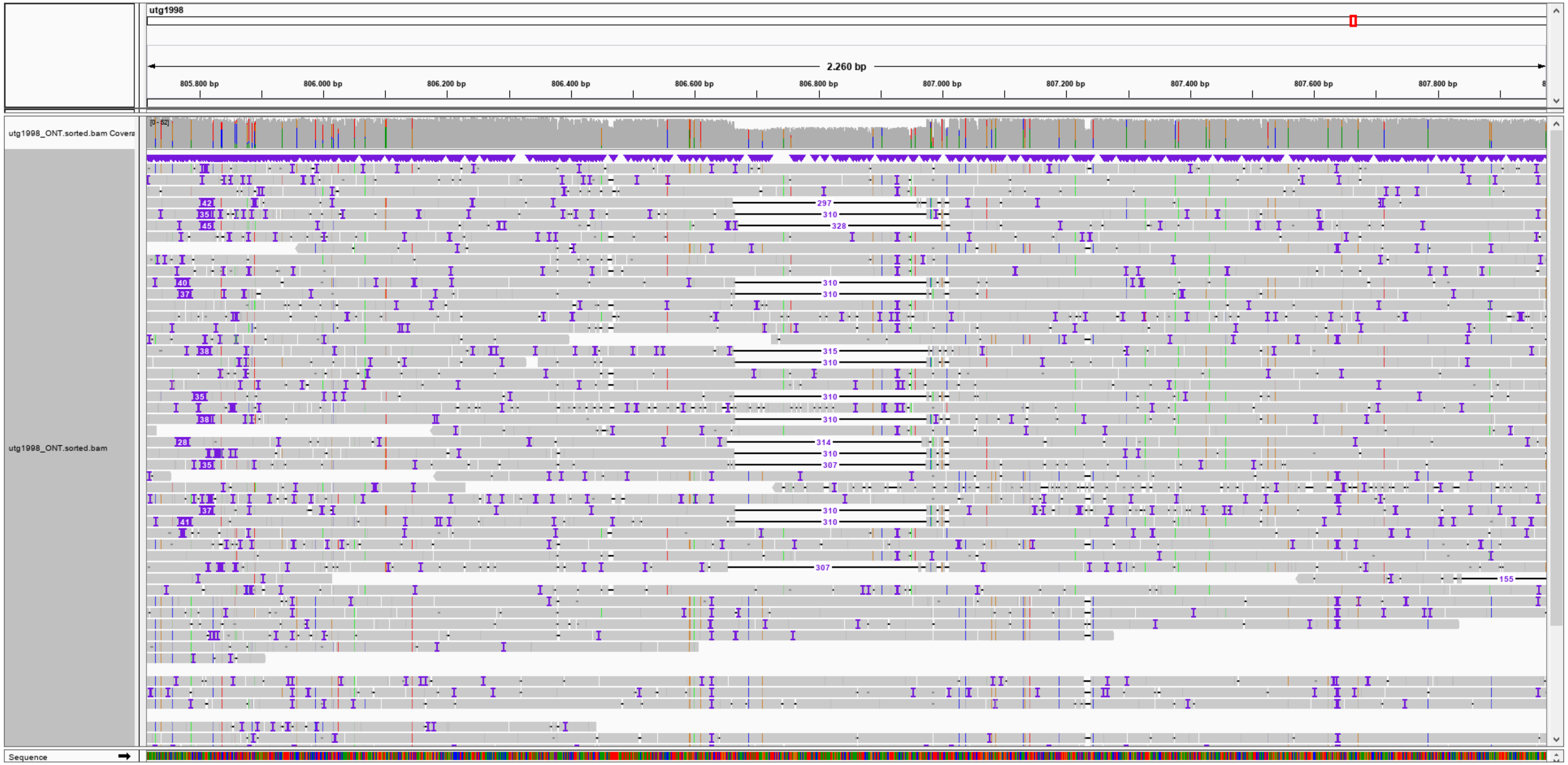


**Overview of marker 807K**

S5 Fig

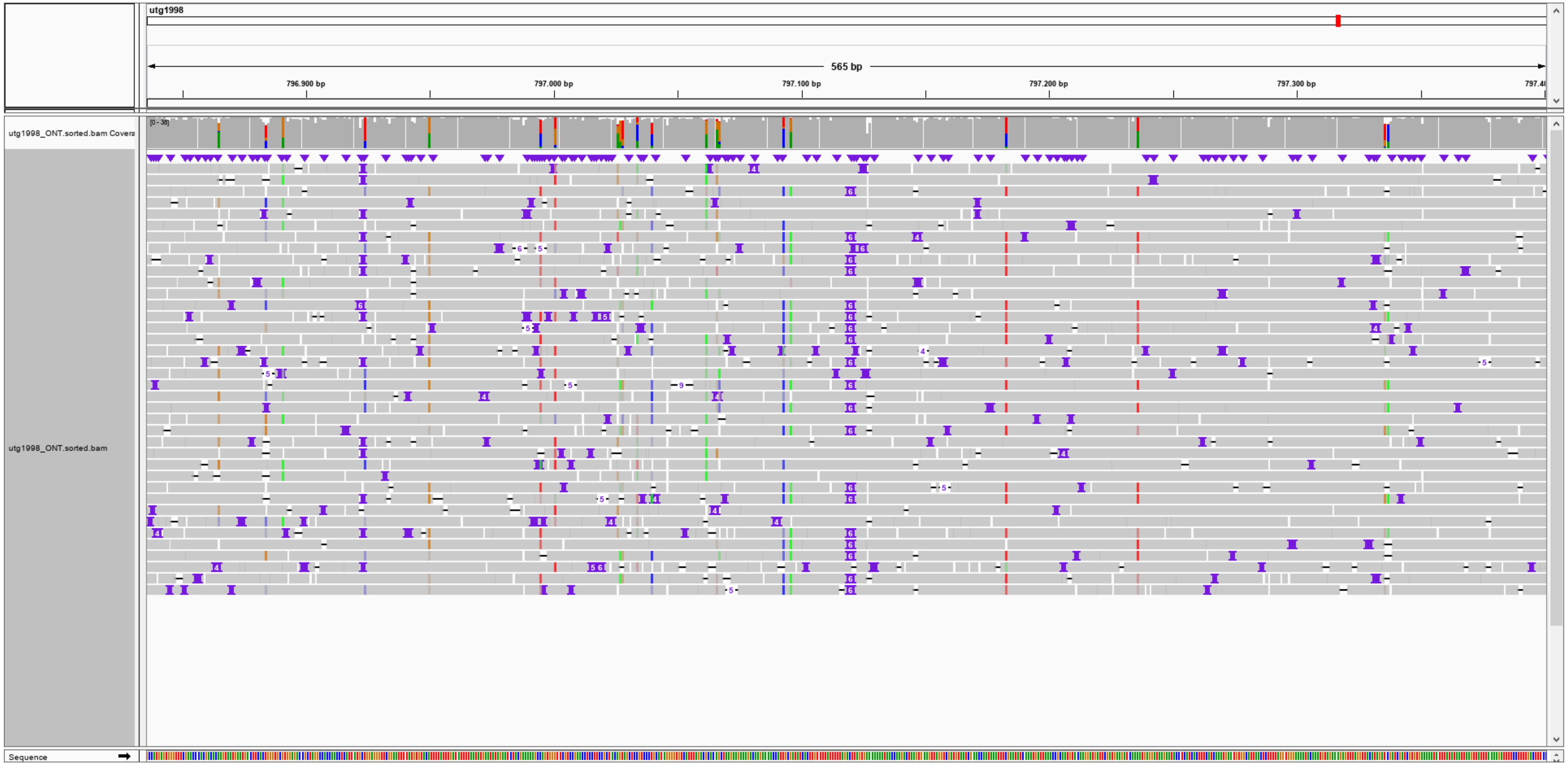


**Overview of marker 797K**

S6 Fig

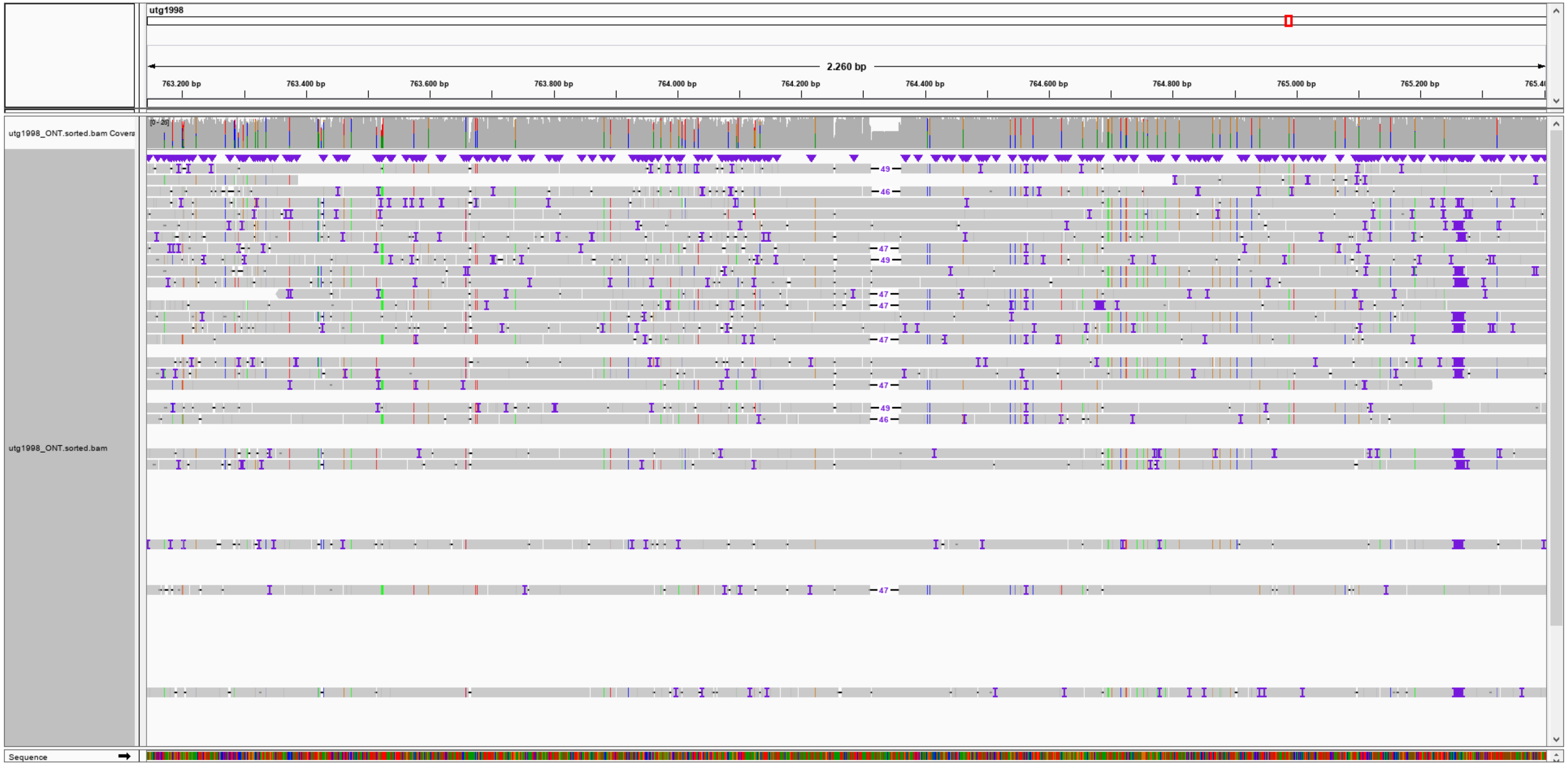


**Overview of marker 764K**
