## Supplementary material for "Tetraose steroidal glycoalkaloids from potato provide resistance against *Alternaria solani* and Colorado potato beetle": S9Fig

S9 Fig

1 … 10 … 20 … 30
1. ScGTR1 MAAMNEVIVA MVPFPDYSHL NQLFVLARFI
2. ScGTR2 MAAINEMIVA MVPFPDYSHL NNLFDLARFI
3. ScGTS MSAINEMIVA MVPFPDYSHL NNLFDLARFI

40 … 50 … 60
1. ScGTR1 ASHNIQVHFL SFVDVNRDLE LRLQGA---S
2. ScGTR2 ASHNIQVHFL CFVDVNRDLK LRLQGSLGVS
3. ScGTS ASHNIQVHFL CFVDVNRDLE LRLQGSLGVS

70 … 80 … 90
1. ScGTR1 NIHFENLLRP SSPTDKDGRD GEPPILMFLE
2. ScGTR2 NIHFENLLRP SSPTDKDGRD GEPPILMFLE
3. ScGTS NIHFENLLRP SSPTDKAGRD GEPPILIFLE

100 … 110 … 120
1. ScGTR1 KLTKPINEIC VQLSTKARKL VIIHDLIMSE
2. ScGTR2 KLTKPINEIC VQLSTKARKL VIIHDLIMSE
3. ScGTS KLTKPINEIC VQLSTKARKL VIIHDLIMSG

130 … 140 … 150
1. ScGTR1 QVTDLNTLLN VKSYMFHTGS TFSRYSSLKQ
2. ScGTR2 QITDLNTLLN VKSYMFHTGS TFSRYSSLKQ
3. ScGTS QVTDLNTLLN IKSYMFHTGS TFSRYSSLKK

160 … 170 … 180
1. ScGTR1 TIPDIVDDDD DHVKLIKQMQ DEFPLLEPHE
2. ScGTR2 TIPDIVDDDD DHVKLIKQMQ DEFPLLEPHE
3. ScGTS LFRT*

190 … 200 … 210
1. ScGTR1 KYCFQVELDD DPVMHYSGEI INSCREMEGK
2. ScGTR2 KYCFQMELDD D--MRYSGEI INSYREMEGK
3. ScGTS

220 … 230 … 240
1. ScGTR1 YPDLLAKAKG KPLWAFGPFH MLLESHSNSS
2. ScGTR2 YPDLLAKAKG KPLWAFGPFH MLLESHSNSS
3. ScGTS

250 … 260 … 270
1. ScGTR1 SNITRHDQCL EFLDKQDVNS VIFVSFGSTT
2. ScGTR2 RNITRHDQCL EFLDKQDVNS VIFVSFGSTT
3. ScGTS

280 … 290 … 300
1. ScGTR1 TLSQDQVNEL ALGLEQSNHK FIWVLRKGDN
2. ScGTR2 TLSQKQVNEL ALGLEQSNHK FIWVLRKGDN
3. ScGTS

310 … 320 … 330
1. ScGTR1 AEKLKEKDIK IELPEGFEER VEGRGIMVNW
2. ScGTR2 AEKLKEKDIK IELPEGFEER VEGRGIMVNW
3. ScGTS

340 … 350 … 360
1. ScGTR1 APQLEILGHS STGGFMSHCG WNSCIESISM
2. ScGTR2 APQLEILGHS STGGFMSHCG WNSCIESISM
3. ScGTS

370 … 380 … 390
1. ScGTR1 GVPVATWPIS YDQPFNAIFL TNLLKIGISV
2. ScGTR2 GVPVATWPIS YDQPFNAIFL TNLLKIGISV
3. ScGTS

400 … 410 … 420
1. ScGTR1 KSWSHRNELI RASTIEKSVK TLMGTIEGEE
2. ScGTR2 KSWSHRNEVI TASTIEKSVK TLMGTIEGEE
3. ScGTS

430 … 440 … 450
1. ScGTR1 MRRRAVELSK QIKNSVRSGG RARKDMESFI
2. ScGTR2 MRRRAVELSK QIKNSVRSGG RARKDMESFI
3. ScGTS

456
1. ScGTR1 SHIIE *
2. ScGTR2 SHIIE *
3. ScGTS
